# *Dnmt3bas* coordinates transcriptional induction and alternative exon inclusion to promote catalytically active Dnmt3b expression

**DOI:** 10.1101/2022.08.08.503222

**Authors:** Mohd Saleem Dar, Isaiah K Mensah, Ming He, Sarah McGovern, Mark C Hall, Hannah Christian Whitlock, Nina Elise Bippus, Madison Ceminsky, Martin L Emerson, Hern J Tan, Humaira Gowher

## Abstract

During mammalian embryogenesis, DNMT3B activity is critical for the genome-wide establishment of DNA methylation. Using naïve ESC differentiation as a model, we elucidated the mechanism by which lncRNA, *Dnmt3bas,* controls the inducible expression and alternative splicing of *Dnmt3b*. Our data showed that *Dnmt3bas* knockdown increased transcriptional induction and decreased H3K27me3 at Dnmt3b cis-regulatory elements post-differentiation. Notably, transcriptional induction of *Dnmt3b* was accompanied by exon inclusion, switching the major isoform from catalytically inactive *Dnmt3b6* to the active *Dnmt3b1*. While *Dnmt3bas* overexpression attenuated *Dnmt3b* induction, it increased the *Dnmt3b1:Dnmt3b6* ratio. This observation was explained by a specific interaction of *Dnmt3bas* with hnRNPL, which promotes exon inclusion. These data suggest that *Dnmt3bas* coordinates alternative splicing and transcriptional induction of Dnmt3b by facilitating the interaction of hnRNPL and RNA Pol II at the Dnmt3b promoter. This two-pronged mechanism would tightly control DNMT3B activity, ensuring the fidelity and specificity of *de novo* DNA methylation during development.

## Introduction

During mammalian development, epigenetic reprogramming, which involves global erasure and re-establishment of DNA methylation, facilitates the acquisition of epigenetic plasticity and limits the inheritance of acquired epimutations (Greenberg and Bourc’his, 2019). DNA methylation is reset by the *de novo* methylation activity of DNMT3A and DNMT3B DNA methyltransferases. In mice, homozygous knockout of *dnmt3a* and *dnmt3b* is embryonic lethal, demonstrating their essential role in mammalian development (Okano et al., 1999; Okano et al., 1998). *Dnmt3b* is dynamically expressed during development, with the highest expression in the early stages. *Dnmt3b^-/-^* ESC (murine embryonic stem cells) show defective differentiation potential and loss of DNA methylation at the regulatory elements of lineage-specific genes and minor satellite repeats, which are the preferred targets of the *Dnmt3b* enzyme (Gopalakrishnan et al., 2009; Norvil et al., 2020; Okano et al., 1999). In somatic cells, the expression of *Dnmt3b* is low and is controlled in a tissue-specific manner (Robertson et al., 2000; Robertson et al., 1999). *Dnmt3b* is transcribed in more than 30 alternatively spliced isoforms, although only a few have been detected at the protein level (Ostler et al., 2007; Xie et al., 1999). Several of these isoforms are catalytically inactive due to the loss of key catalytic residues. The two major isoforms expressed in normal cells are *Dnmt3b1* and *Dnmt3b6*. The exclusion of exons 22 and 23 results in the loss of a significant part of the target recognition region in the Dnmt3b6 transcript, thus rendering the protein enzymatically inactive. However, DNMT3B6 has been shown to interact and allosterically activate the full-length catalytically active enzyme DNMT3B1 (Duymich et al., 2016; Ostler et al., 2007; Saito et al., 2002; Wang et al., 2007; Weisenberger et al., 2004; Zeng et al., 2020).

The aberrant increase in *DNMT3B* expression leads to DNA hypermethylation and loss of gene regulation in various human diseases (Esteller, 2005; Walton et al., 2014; Watanabe and Maekawa, 2010). Sequence polymorphisms in the *DNMT3B* promoter that increase promoter activity are associated with an increased risk of several cancers (Montgomery et al., 2004; Shen et al., 2002; Singal et al., 2005). Loss of function mutations in human *DNMT3B* are highly prevalent in patients with ICF (immunodeficiency, centromeric instabilities, and facial abnormalities) syndrome and cause hypomethylation of repetitive elements and genomic instability in B cells (Gowher and Jeltsch, 2002; Okano et al., 1999; Xie et al., 2006; Xu et al., 1999). Aberrant expression and splicing of *DNMT3B* are linked to the loss of methylation at oncogenes and repetitive elements in diverse cancers, including colorectal, lung, and breast cancers (Guil and Esteller, 2009; Kulis and Esteller, 2010; Mensah et al., 2021; Szyf, 2005). These observations indicate that the highly regulated spatiotemporal expression and alternative splicing of *DNMT3B* are critical for cell differentiation, homeostasis, and survival (Huntriss et al., 2004; Okano et al., 1998). Despite the breadth of evidence supporting the critical role of *DNMT3B* in differentiation and cell identity, little is known about the mechanisms that control its transcription.

The murine *Dnmt3b* gene locus comprises a CpG island promoter and an enhancer element ∼8 kb upstream of the *Dnmt3b* promoter (Ishida et al., 2003; Jinawath et al., 2005). A divergently expressed antisense transcript (*Dnmt3bas*) initiates 1 kb downstream of the *Dnmt3b* promoter and is potentially a long non-coding RNA (lncRNA). LncRNAs have critical cellular functions, including transcription, splicing, translation, RNA export, chromatin looping, and DNA repair. Most lncRNAs are transcribed by RNA Pol II but at lower levels, and many undergo capping, splicing, and polyadenylation (Mele et al., 2017; Statello et al., 2021; Zuckerman and Ulitsky, 2019). LncRNAs interact with chromatin proteins and regulate the activation and silencing of genes (Derrien et al., 2012; Guo et al., 2020; Guttman et al., 2009; Guttman et al., 2011; Tian and Manley, 2017). During ESC differentiation, several lncRNAs originate from the divergent transcription. These lncRNA/mRNA gene pairs are coordinately regulated and required for pluripotency and differentiation (Guttman et al., 2011; Luo et al., 2016). Here we report *Dnmt3bas/Dnmt3b* as a coordinately expressed gene pair where the lncRNA*, Dnmt3bas,* regulates transcriptional induction and alternative splicing of *Dnmt3b* mRNA.

During differentiation of naive pluripotent mouse embryonic stem cells (2i-ESCs), the temporal expression pattern of *Dnmt3b* is similar to that observed during embryonic development (Ficz et al., 2013; Leitch et al., 2013; Ying et al., 2008), making them an ideal model system. Here we show that in undifferentiated 2i-ESCs, *Dnmt3b* is expressed at a low basal level and primarily as *Dnmt3b6*, the catalytically inactive isoform. The expression of Dnmt3b is strongly induced in response to the differentiation signal and is, interestingly, accompanied by exon inclusion, which promotes the expression of the catalytically active, *Dnmt3b1* as the major isoform. As differentiation proceeds, *Dnmt3b* expression is downregulated. Our study is the first to report the role and mechanism of *Dnmt3bas* in regulating *Dnmt3b* expression and alternative splicing.

Gene expression analysis of undifferentiated 2iESCs and cells post differentiation showed a coordinated yet contrasting expression pattern of *Dnmt3b* and *Dnmt3bas*, with the *Dnmt3b* pattern mimicking the expression of *Dnmt3b* observed *in vivo*. The chromatin modification state of proximal and distal enhancers and enhancer-promoter looping complemented the *Dnmt3b* transcriptional state. Whereas we observed no discernable effect of the downregulation (KD) or overexpression (OE) of *Dnmt3bas* on *Dnmt3b* basal expression, the transcriptional induction of *Dnmt3b* was decreased in *Dnmt3bas* OE cells and increased in *Dnmt3bas* KD cells. Interestingly, the undifferentiated OE cells showed an increase in H3K27me3 at the cis-regulatory elements, suggesting the role of *Dnmt3bas* in regulating PRC2 activity at these sites. Additionally, higher enrichment of H3K27Ac and an increased interaction frequency between enhancer and promoter were observed in the KD cells post-differentiation. In *Dnmt3bas* OE cells, we also observed increased exon inclusion, resulting in higher *Dnmt3b1:Dnmt3b6* ratio. Our systematic experimental analysis determined the splicing factor, hnRNPL, as the binding partner of *Dnmt3bas* that facilitates exon inclusion during transcriptional induction of *Dnmt3b*.

Overall, this comprehensive study elucidated a mechanism by which a promoter-associated lncRNA, *Dnmt3bas*, coordinates transcriptional induction and alternative splicing of an essential developmental gene and establishes the role of cis-regulatory elements in this process.

## Results

### Transcriptional induction and alternative splicing of Dnmt3b

We adapted serum-cultured murine embryonic stem cells (s-ESCs) to 2i media (2i-ESCs). As reported earlier, naïve pluripotency markers were upregulated during the adaptation process. Moreover, the expression of *Dnmt3a* and *Dnmt3b* were downregulated, with the concomitant loss of DNA methylation genome-wide (Figure S1A-1C). We differentiated 2i-ESCs and collected samples from D1-D6 post-differentiation (Fig S1D). Differentiation was monitored by a change in the expression of pluripotency and differentiation markers by RT-qPCR (Fig S1E). Analysis of *Dnmt3b* expression showed a 10-15 fold increase, which peaks at D3, followed by a substantial decrease that stays steady from D4 to D6 post-differentiation (Figure 1A). Since *Dnmt3b* is expressed at a low basal level in 2i-ESCs, the induction is significantly higher than observed during the differentiation of s-ESCs (Figure S1F), making 2i-ESC differentiation an ideal experimental model system.

**Figure 1.**
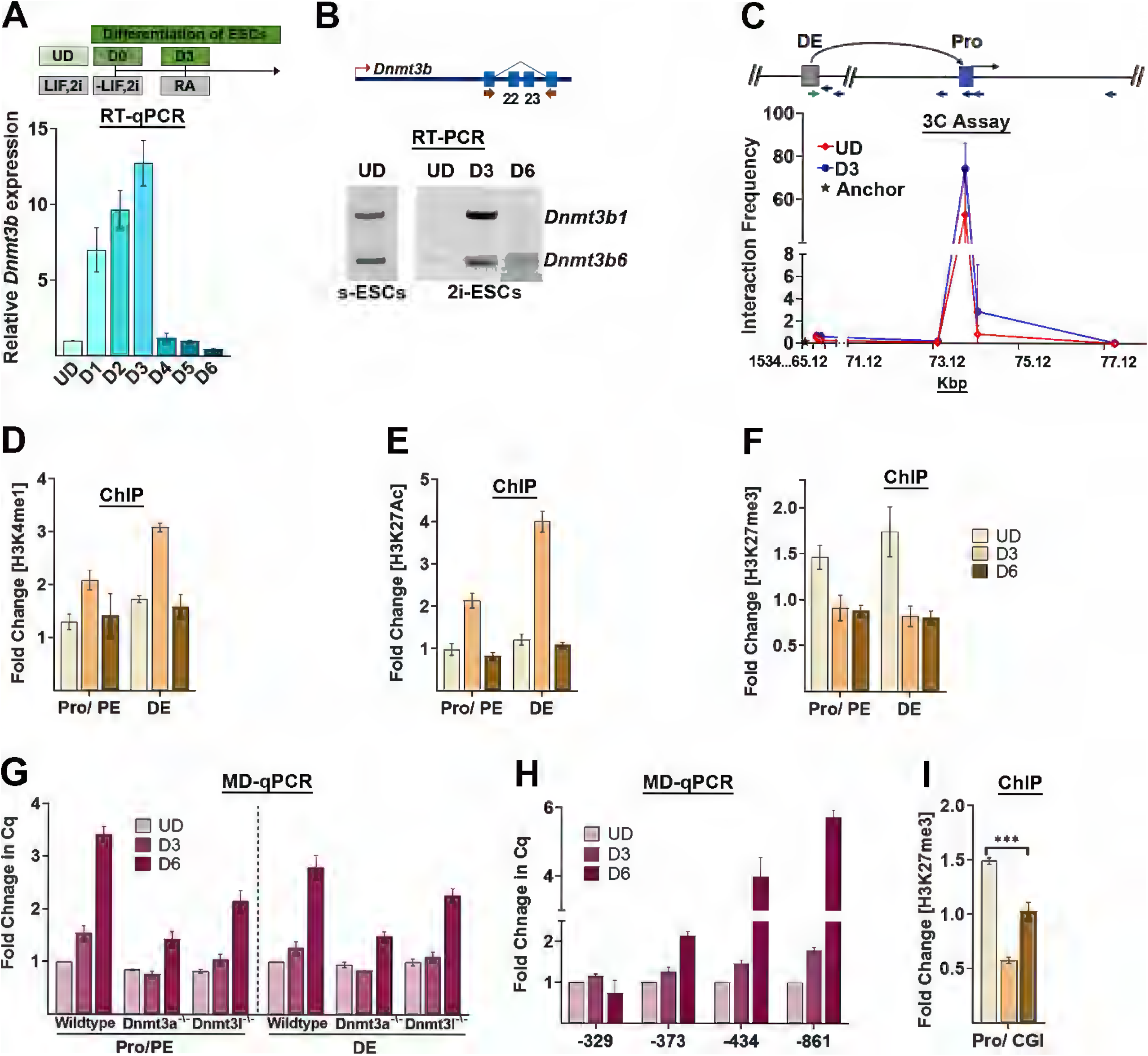
Proximal and distal enhancers regulate Dnmt3b induction. UD: undifferentiated; D1-D6: Days post-induction of differentiation; ESCs: embryonic stem cells; s-ESC: serum cultured ESCs; RA: Retinoic Acid; LIF: Leukocyte Inhibitory factor; 2i: Signaling pathway Inhibitors, CHIR99021 and PD184352; PE: proximal enhancer; Pro: promoter; DE: Distal enhancer, CGI: CpG island; WT: wild type; 3a: Dnmt3a; 3l: Dnmt3L (A) RT-qPCR of Dnmt3b during 2iESC differentiation as illustrated. The threshold cycle (Ct) values were normalized to Gapdh, and expression is shown relative to that in UD cells. The data show a steady increase in Dnmt3b expression up to D3, which decreases at D4 post-differentiation. (B) Illustration of Dnmt3b alternative exons, 22 and 23, and position of the primers used to amplify the alternatively spliced isoforms, *Dnmt3b1* and *Dnmt3b6*. RT-PCR analysis *Dnmt3b* alternative splicing in serum cultured (S-ESCs) and 2i cultured (2i-ESCs) cells pre and post-differentiation. (C) Chromatin conformation assay (3C) shows an interaction of the distal enhancer with the promoter of Dnmt3b. The location of the distal enhancer and promoter in the Dnmt3b locus, the 3C PCR primers, and the corresponding HaeIII sites (tick marks) are shown. qPCR analysis was done using the primer at the DE as an anchor (Chr2 position: 153466046), and interactions with downstream sites were measured in 2iESC at undifferentiated and D3 post differentiation state. Plotted are the relative interactions in arbitrary units of the Dnmt3b DE with six HaeIII sites. The X-axis represents the chromosome 2 location (153465120-153477120) of these HaeIII sites. A significant increase in the interaction frequency was observed between DE and Pro of the Dnmt3b gene at D3 post-differentiation. (D, E, F, I) Chromatin immunoprecipitation (ChIP)-qPCR shows fold enrichment over input. Histone modifications at Dnmt3b regulatory elements (D) H3K4me1 (E) H3K27Ac and (F, I) H3K27me3 in 2iESCs pre- and post-differentiation. Concomitant with Dnmt3b induction, a systematic increase and decrease in the H3K4me1 and H3K27Ac signal was observed post-differentiation. Whereas H3K37me3 demethylation is maintained post differentiation at the Pro/PE and DE, it is regained at the CGI region of the Pro, close to the TSS of the Dnmt3b gene. P-values were derived from Student’s t-test: *p < 0.05; **p < 0.01; ***p < 0.005. (G, H) MD-qPCR was used to measure DNA methylation. The DNA was restricted using the methylation-dependent enzyme MspJI which cuts at methylated cytosines. The specified regions in the regulatory elements were amplified using the cleaved DNA as the template by qPCR. The Cq values for differentiated samples were normalized to that of the undifferentiated sample. An increase in the Cq value indicates a gain in DNA methylation. (G) Genomic DNA was collected from the WT, Dnmt3a^-/-^ and Dnmt3l^-/-^ ESC cell lines pre-and post-differentiation. DNA methylation analysis shows a reduced gain in both Dnmt3a^-/-^ and Dnmt3l^-/-^ cells. (H) DNA methylation analysis in 2iESCs pre and post-differentiation at Dnmt3b Pro and CGI regions. The tested CpG sites are represented on X-axis as base pairs upstream of the Dnmt3b TSS. The data show a steady gain of DNA methylation corresponding to the distance away from the CGI, which is maintained in an unmethylated state. Results are presented as normalized mean values ± SEM for n = 3. See also Figure S1.

*Dnmt3b* transcripts comprise two major alternatively spliced isoforms, *Dnmt3b1*, a full-length transcript, and *Dnmt3b6*, in which exons 22 and 23 are excluded (Figure 1B). Whereas *Dnmt3b1* codes for an active DNA methyltransferase enzyme, the *Dnmt3b6* protein is inactive due to the absence of motifs critical for catalytic activity (Duymich et al., 2016; Ostler et al., 2007; Saito et al., 2002; Wang et al., 2007; Weisenberger et al., 2004; Zeng et al., 2020). RT-PCR was used to amplify both isoforms pre- and post-differentiation of ESCs (Figure 1B). As shown previously (Gowher et al., 2008), both isoforms are expressed equally in FBS-cultured ESCs. Interestingly, in 2i-cultured ESC, *Dnmt3b* mainly comprises the shorter isoform *Dnmt3b6*, indicating that exon exclusion is preferred during basal transcription of *Dnmt3b*. However, upon differentiation, as *Dnmt3b* expression is induced, *Dnmt3b1* is expressed as the major isoform indicating that the alternative splicing switches in favor of exon inclusion during transcriptional induction Post-differentiation transcriptional repression is again accompanied by switching to exon exclusion, and *Dnmt3b6* is expressed as a major isoform in embryoid bodies (Figure 1B, S1G). These data suggest that in conjunction with transcriptional induction, the alternative exon inclusion/exclusion regulates the level of catalytically active DNMT3B, underscoring the critical role of the alternative splicing.

The cis-regulatory elements in the Dnmt3b locus constitute a CpG island promoter and putative proximal and distal enhancer elements located about 0.3 and 8 kilobases upstream of the transcription start site (TSS), respectively (Figure S1H) (ENCODE CRE) (Ficz et al.). To determine the engagement of the putative distal enhancer in the regulation of *Dnmt3b* expression, we examined the enhancer-promoter (E-P) looping interaction pre- and post-differentiation in 2i-ESCs, using a chromatin conformation capture (3C) assay (Hagege et al., 2007; Naumova et al., 2012). We observed a potent and specific contact between the distal enhancer and promoter regions. The E-P loop was prevalent in the undifferentiated 2i-ESC when *Dnmt3b* was expressed at the basal level. However, a significant increase in the interaction frequency was observed on D3 post-differentiation. These observations suggest that pre-positioning the enhancer next to the promoter enables a quick transcriptional response to the induction signal (Figure 1C).

Based on the transcriptional state of the associated gene, enhancer regions acquire different chromatin modifications. We measured the enrichment of H3K4me1, H3K27Ac, and H3K27me3 at *Dnmt3b* regulatory regions pre- and post-differentiation. At D3 post-differentiation, we observed an expected increase in H3K4me1 and H3K27Ac concomitant with enhancer activation and induction of *Dnmt3b* expression. *Dnmt3b* repression at D6 post-differentiation was associated with a decrease in H3K4me1 and H3K27Ac at both proximal and distal enhancer regions (Figures 1D and 1E). In contrast, H3K27me3 at the proximal and distal enhancer regions was higher in undifferentiated 2iESCs (UD), followed by a decrease post-differentiation at D3 and D6 (Figure 1F). It is noteworthy that deacetylation of H3K27 at D6 post-differentiation is not succeeded by H3K27 methylation at these sites. H3 occupancy shows no significant difference in enrichment in various samples pre- and post-differentiation, further validating the significance of the observed chromatin modification changes (Figure S1I).

Previous studies have shown that during enhancer silencing, histone demethylation of H3K4me1 and deacetylation of H3K27Ac by Lsd1-Mi2NURD-complex poises the chromatin for DNMT3A activity (Petell et al., 2016). Therefore, we asked if DNA methylation at enhancer regions maintained the repression of *Dnmt3b* post-differentiation. Methylation-dependent qPCR (MD-qPCR) showed a significant gain of DNA methylation at distal and proximal enhancer regions on D6 post-differentiation (Figure 1G). To directly test the role of DNMT3A and DNMT3L, we compared DNA methylation in the WT, *Dnmt3a^-/-,^* and *Dnmt3l^-/-^* cells. We observed that compared to WT and *Dnmt3l^-/-^* cells, the *Dnmt3a^-/-^* cells showed a severe defect in gaining DNA methylation (Figure 1G). The *Dnmt3b* promoter also harbors a CpG island (CGI) around TSS, and we asked if DNA methylation spreads into the CGI. As previously reported for strong CGIs, the DNA methylation levels decreased with increasing proximity to the *Dnmt3b* promoter CGI (Figure 1H). However, unlike in the proximal and distal enhancers regions (Fig 1D), the CGI promoter region showed a recovery of H3K27me3 at D6 post-differentiation, demonstrating that the PRC2 - mediated mechanism maintains the repressed state of the *Dnmt3b* promoter (Figure 1I). Overall, these data establish the activity of distal and proximal enhancers in regulating *Dnmt3b* expression during 2i-ESC differentiation.

### Dnmt3bas regulates the magnitude of Dnmt3b induction

The divergent antisense (*as*) long non-coding RNA (lncRNA), *Dnmt3bas*, is a 2.8 kb transcript that initiates 800 bps downstream of TSS in the *Dnmt3b* promoter (Figure S2A). Sequence analysis of *Dnmt3bas* revealed a potential exon-intron structure, and structural modeling of the spliced *Dnmt3bas* using RNA Fold webserver (ViennaRNA) showed a stem structure typical of lncRNAs (Figure S2B). Based on the predicted exon-intron boundaries, we designed primers to capture potential spliced and polyadenylated forms of *Dnmt3bas* (Figure S2A). An expected unique band of nearly 716 bps, which corresponds to the size of spliced *Dnmt3bas*, confirmed the splicing and polyadenylation of *Dnmt3bas* (Figure 2A). Subcellular localization assays showed that *Dnmt3bas* is predominantly present in the nucleus (Figure 2B).

**Figure 2.**
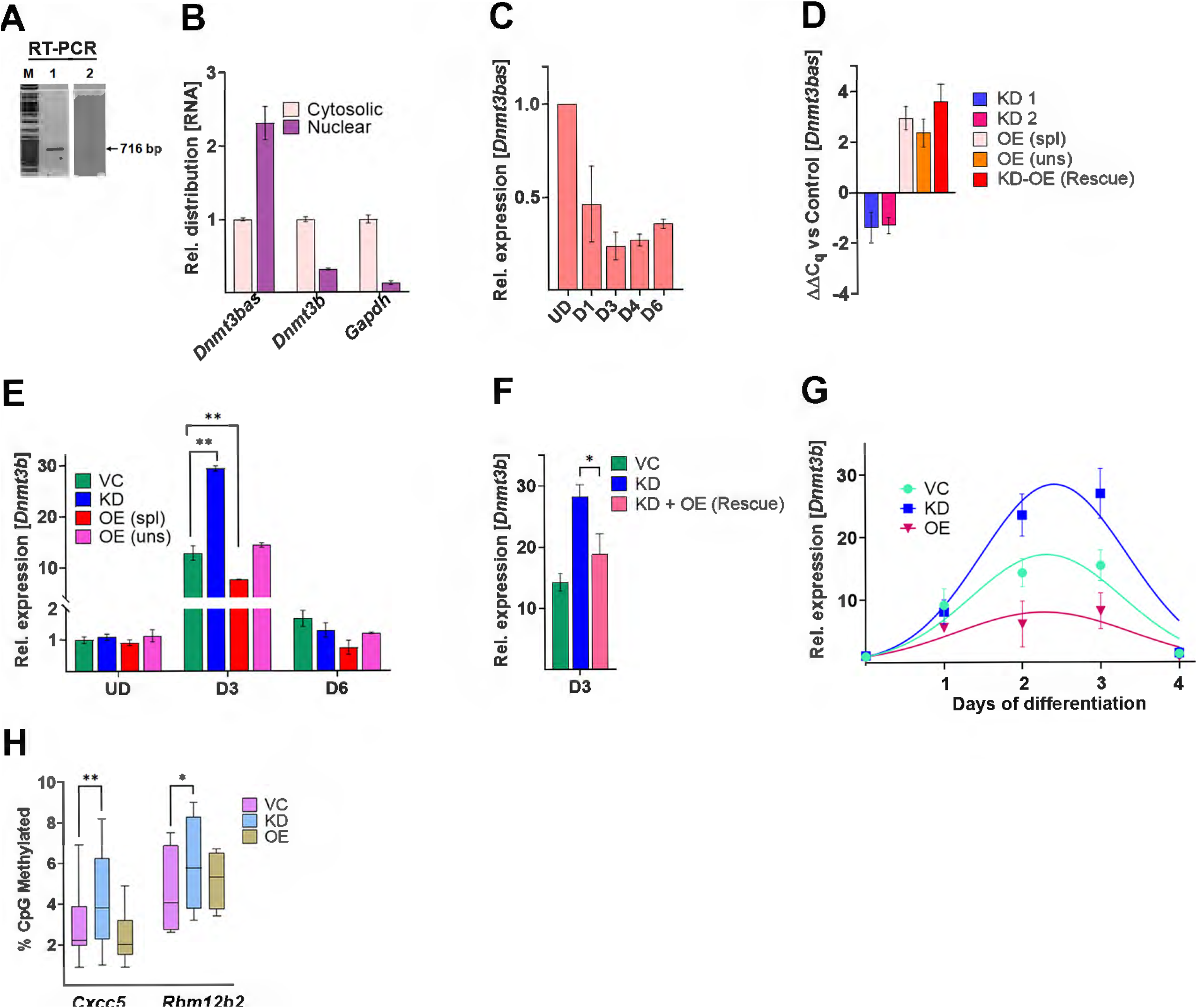
Spatiotemporal expression of Dnmt3b in *Dnmt3bas* manipulated ESCs. UD: undifferentiated; D1-D6: Days post-induction of differentiation; ESCs: embryonic stem cells; VC: Vector control; KD: shRNA mediated knockdown of *Dnmt3bas*; OE: overexpression of *Dnmt3bas* lncRNA; spl: spliced *Dnmt3bas*, uns: unspliced *Dnmt3bas* (A) Ethidium bromide-stained agarose gel. RT-PCR of *Dnmt3bas* using exonic (lane 1) and Oligo dT (lane 2) primers show a single band corresponding to the spliced transcript. (B, C, D, E, F, G) RT-qPCR analysis of *Dnmt3bas* expression. (B) RT-qPCR of *Dnmt3bas* from nuclear and cytosolic fractions of RNA showed about 2-fold higher enrichment in the nucleus compared to the cytosol. In contrast, Dnmt3b and Gapdh mRNA transcripts showed a significantly higher enrichment in cytosol compared to the nucleus. (C) RT-qPCR shows *Dnmt3bas* expression in 2iESCs pre- and post-differentiation, with the highest expression in the UD state. (D) RT-qPCR of *Dnmt3bas* from transgenic 2i-ESC lines expressing *Dnmt3bas* shRNA (KD1, KD2), spliced (spl) and unspliced (uns) transcripts of *Dnmt3bas* (OE), and spliced *Dnmt3bas* in KD cells (KD-OE). The Ct values were normalized to Gapdh, and expression is shown relative to that in vector control (VC) cells. (E, F) RT-qPCR analysis of Dnmt3b expression in Dnmt3bas manipulated transgenic 2i-ESCs, pre-and post-differentiation. (E) At D3, the KD and OE cells showed significantly higher and lower induction than VC cells, respectively. (F) In KD-OE cells, Dnmt3b induction is rescued to the level in VC cells at D3 post differentiation. (G) Dnmt3b expression in VC, OE, and KD cell lines was measured every 24 hours post-differentiation. The data were fit to a non-linear regression in PRISM to estimate the time of the Dnmt3b induction peak. P values were derived from the ANOVA test or Student’s t-test: *p < 0.05; **p < 0.01; ***p < 0.005. (H) DNA methylation analysis using Bis-Seq. Bisulfite-treated genomic DNA from D2 differentiated cells was used to PCR amplify two Dnmt3b-specific target regions, CXXC5: chr18; 35858578 : 35858863, Rbm12b2; chr4;12113443 : 12114006. The amplicons were sequenced on a high throughput sequencing platform (Wide-Seq), and the data were analyzed using Bismark software. The box plot represents the range of percent DNA methylation at various CpG sites in each target region. The median methylation at both targets is higher in KD cells compared to VC cells. P values were calculated using Wilcoxon -matched pairs rank test: *p < 0.05; **p < 0.01; ***p < 0.005. Results are presented as normalized mean values ± SEM. See also Figure S2.

We first asked if there is a correlation between the expression of *Dnmt3b* and *Dnmt3bas* during 2i medium adaptation and differentiation of ESCs. We used exon-specific primers to detect the expression of spliced *Dnmt3bas*, which also minimizes potential amplification from genomic DNA contamination in the RNA samples (Figure S2C). The RT-qPCR specificity was ensured by the absence of signal in the NRT (no reverse transcriptase) control and by visualizing the amplified product as a single band on an agarose gel (Figure S2D). Expression analysis showed that *Dnmt3b* was downregulated at passage P6 and maintained a basal level in 2i-ESCs. Interestingly, *Dnmt3bas* expression constantly increased during the adaptation process (Figure S2E). However, during 2i-ESC differentiation, the expression of *Dnmt3bas* was downregulated concomitant with induction of the *Dnmt3b* (Figure 2C).

The coordinated yet contrasting expression pattern of the *Dnmt3b*/*Dnmt3bas* pair suggests a potential role for *Dnmt3bas* in regulating *Dnmt3b* transcription. Therefore, we generated stable ESCs lines expressing anti-*Dnmt3bas* shRNA (KD1 and KD2) or overexpressing (OE) spliced and unspliced forms (spl and uns) of *Dnmt3bas* and successfully adapted them to 2i medium shown by an increase in *Prdm14* expression in adapted cells (Figure S2F, G). Gene expression analysis in the *Dnmt3bas* KD1 and KD2 cells showed a significant reduction of *Dnmt3bas* by nearly 2.5-fold compared to the vector control (VC) cells. Conversely, *Dnmt3bas* OE cells showed nearly 8-9-fold higher expression of *Dnmt3bas* compared to VC cells (Figure 2D). Furthermore, cell fractionation analysis of the *Dnmt3bas* KD and OE cells showed a proportional change in the *Dnmt3bas* transcript level in the cytoplasm and nucleus (Figure S2H), supporting the localization of *Dnmt3bas* primarily in the nucleus. We also ruled out the potential confounding impact of impaired pluripotency in the *Dnmt3bas* manipulated cell lines. Cell morphology and gene expression analysis pre and post-differentiation show no significant difference between the three cell lines (Figure S2I, J).

Next, we determined the effect of *Dnmt3bas* manipulations on *Dnmt3b* expression in the *Dnmt3bas* KD and OE cell lines. The data showed no difference in *Dnmt3b* basal expression in undifferentiated 2i-ESCs (Figure 2E). We next tested the effect on *Dnmt3b* induction post-differentiation. Similar to our observation in Figure 1A, the VC cells showed a 15-fold induction of *Dnmt3b* at D3 before reaching the basal level at D6 post-differentiation. Interestingly, in the KD cells, the induction of *Dnmt3b* at D3 increased nearly 30-fold and returned to the basal level at D6 post-differentiation. In OE cells, an opposite effect on *Dnmt3b* induction was observed, with only a 7-fold increase in *Dnmt3b* expression on D3 post-differentiation. Moreover, the repressive activity of *Dnmt3bas* was specific for only the spliced isoform since cells overexpressing the unspliced *Dnmt3bas* behaved similarly to VC cells (Figure 2E). In addition, this effect on *Dnmt3b* induction was rescued by overexpression of *Dnmt3bas* in KD cells (KD-OE), confirming the trans activity of the *Dnmt3bas* transcript (Figure 2F). A similar effect on Dnmt3b expression was observed in both KD1 and KD2 cells (Figure S2K); therefore, only KD1 cells were used for further investigation. We next asked if *Dnmt3bas* modulated the kinetics or magnitude or both of *Dnmt3b* induction. A time-course experiment was performed to measure the temporal increase in transcript levels of *Dnmt3b* every 24 hrs post differentiation. The data were fit to a non-linear regression using GraphPad Prism. Whereas an apparent change in the magnitude of induction was recorded, it peaked at the same time point in all three cell lines, indicating the effect of *Dnmt3bas* on the magnitude of *Dnmt3b* induction (Figure 2G).

2iESC are hypomethylated, and genome-wide establishment of DNA methylation requires activities of both DNMT3A and DNMT3B post-differentiation (Okano et al., 1999). We investigated the potential impact of *Dnmt3bas* manipulations on the establishment of DNA methylation. Analysis of global methylation using methylation-sensitive restriction showed a delay in the gain of methylation in both KD and OE cells compared to VC cells (Figure S2L). We used bisulfite sequencing to compare gain in DNA methylation of two Dnmt3b-specific target regions on D2 post-differentiation (Yagi et al., 2020) (Figure 2H). A significant increase in DNA methylation was observed in KD cells compared to the VC and OE cells, demonstrating the effect of higher *Dnmt3b* expression on DNMT3B target methylation.

Collectively, these data confirm the role of the spliced *Dnmt3bas* transcript in *Dnmt3b* transcriptional induction and suggest a mechanistic model in which the repressive effect of *Dnmt3bas* maintains *Dnmt3b* promoter/proximal enhancer in the primed state in undifferentiated cells to fine-tune the magnitude of *Dnmt3b* induction in response to differentiation signals.

### Dnmt3bas modulates the chromatin modification state at Dnmt3b regulatory regions

To understand the mechanism of *Dnmt3bas* in regulating *Dnmt3b* induction, we measured chromatin modifications, i.e., H3K4me1, H3K27me3, and H3K27Ac at *Dnmt3b* regulatory regions during differentiation and compared changes in KD and OE cells to VC cells. In all three cell lines, an increase in H3K4me1 was observed at both proximal and distal enhancer regions at D3 post-differentiation compared to the undifferentiated (UD) cells (Figure 3A). Similarly, the loss of H3K27me3 was accompanied by the gain of H3K27Ac at the proximal and distal enhancer region in D3 cells (Figure 3B). Interestingly, we observed a lower enrichment of H3K27Ac on D3 in OE cells than in KD and VC cells (Figure 3C), corresponding to the alleviated induction of *Dnmt3b* gene expression (Figure 2F).

**Figure 3.**
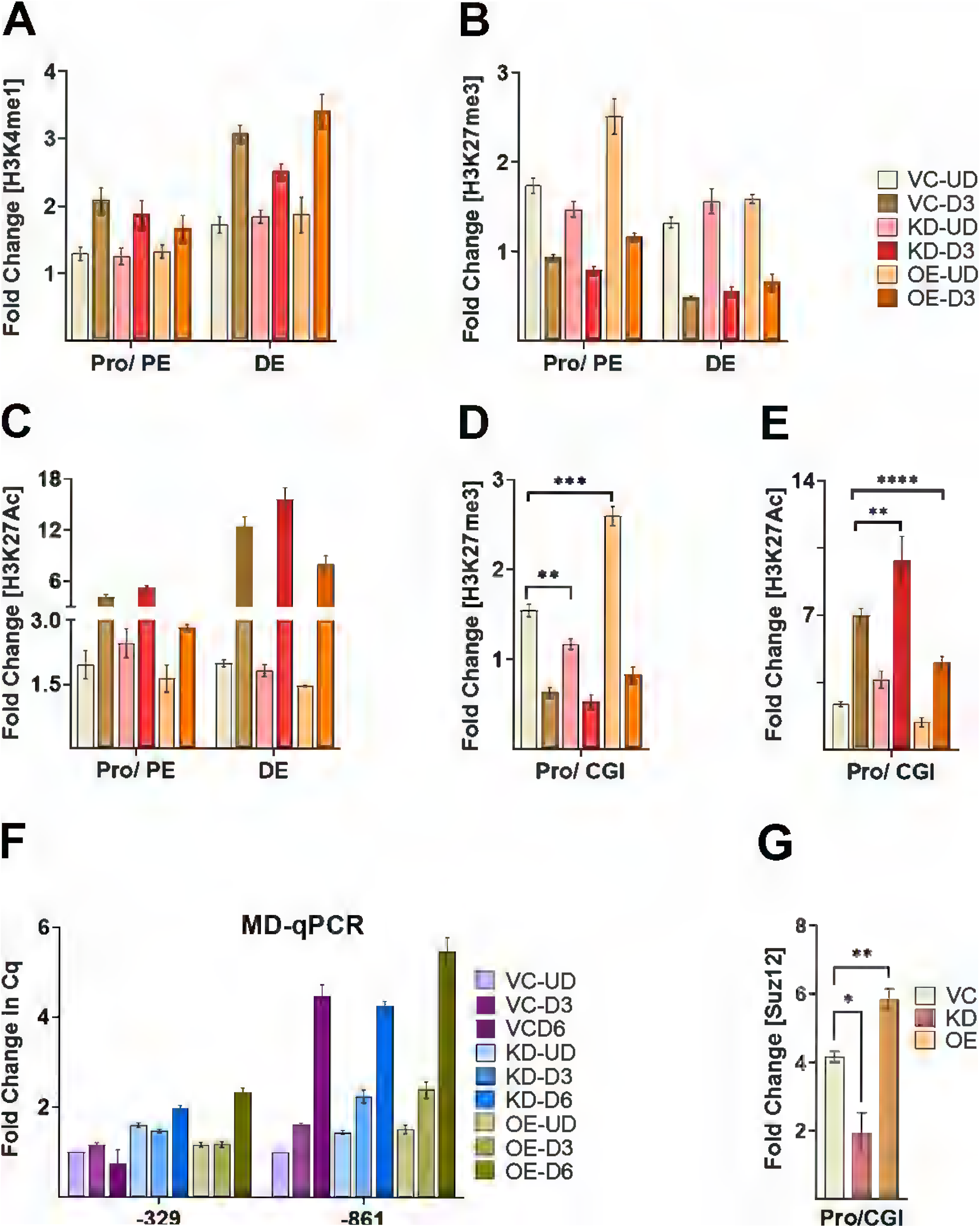
Chromatin modification at Dnmt3b regulatory elements in *Dnmt3bas* manipulated ESCs. UD: undifferentiated; D3, D6: Days post-induction of differentiation; PE: proximal enhancer; Pro: promoter; DE: Distal enhancer; CGI: CpG island; VC: Vector control; KD: shRNA mediated knockdown of *Dnmt3bas*; OE: overexpression of *Dnmt3bas* lncRNA (A, B, C, D, E, G) ChIP-qPCR assays show fold enrichment over input normalized to control. Histone modifications at Dnmt3b regulatory elements (A) H3K4me1 (B, D) H3K27me3 and (C, E) H3K27Ac pre- and D3 post-differentiation in VC, KD, and OE cells. (A) On D3 post-differentiation, an increase in the H3K4me1 signal was observed in KD and OE cells, similar to the VC cells in the Pro/PE and DE regions. (B, D) All cell lines show H3K27me3 demethylation on D3 post-differentiation. However, a higher enrichment of H3K27me3 is observed in undifferentiated (UD) OE cells at the Dnmt3b promoter, including PE and CGI regions. (C, E) H3K27Ac shows an increase at D3 post differentiation, albeit less in OE cells at all the cis-regulatory elements. A significantly higher enrichment in KD cells compared to VC cells is also notable at the CGI region on D3 post-differentiation. (G) Fold enrichment of the PRC2 complex component, Suz12, at the *Dnmt3b* promoter CGI region shows an opposite effect of *Dnm3bas* OE and KD on Suz12 binding. (F) DNA methylation at the Dnmt3b promoter was measured by MD-qPCR. After digestion of genomic DNA from VC, KD, and OE cells pre and post-differentiation using the FspE1 enzyme, the specified regions in the regulatory elements were amplified using qPCR. The Cq values for differentiated samples were normalized to that of the VC-UD sample. An increase in the Cq value indicates a gain in DNA methylation. The tested CpG sites are represented on X-axis as base pairs upstream of the Dnmt3b TSS. The data show a steady gain of DNA methylation corresponding to the distance away from the CGI, which is maintained in an unmethylated state. p values were derived from ANOVA test: *p < 0.05; **p < 0.01; ***p < 0.005. Results are presented as normalized mean values ± SEM. See also Figure S3.

We next measured H3K27me3 and H3K27Ac close to CGI at the Dnmt3b promoter. Our data showed that H3K27me3 demethylation is accompanied by H3K27 acetylation during the differentiation of all three cell lines (Figure 3D, 3E). However, compared to the VC cells, we observed a lower and a higher enrichment of H3K27me3 in undifferentiated KD and OE cells, respectively (Figure 3D, S3A). Expectedly, an opposite trend in H3K27Ac was observed in these cells (Figure 3E), conforming to the difference in Dnmt3b induction. The absence of predicted changes in H3K27me3 at the distal enhancer in *Dnmt3bas* KD and OE cells is possibly due to the low enrichment of this modification, limiting its detection. We also tested the potential effect of *Dnmt3bas* on DNA methylation at the promoter and proximal enhancer (PE) region. Similar to the VC cells, a gain of DNA methylation was observed in promoter regions farther from the CGI that was not affected by changes in *Dnmt3bas* levels in KD and OE cells (Figure 3F).

Based on the above data, we speculated that *Dnmt3bas* recruits the PRC2 complex to *Dnmt3b* cis-regulatory regions, similar to several other lncRNAs known to escort the PRC2 complex to their target sites (listed in (Schertzer et al., 2019; Statello et al., 2021)). The presence of PRC2 complex was confirmed by Suz12 ChIP, which shows strong enrichment at the Dnmt3b promoter. In alignment with H3K27me3, significantly lower and higher Suz12 enrichment was observed in the KD and OE cells, respectively, compared to VC cells (Figure 3G). This observation suggests the role of *Dnmt3bas* in recruiting the PRC2 complex. To determine the interaction of the PRC2 complex with *Dnmt3bas*, we performed an RNA pull-down assay using *in vitro* transcribed biotinylated *Dnmt3bas* and immunoblotted the precipitate for the PRC2 complex proteins. An absence of a positive signal suggested either weak direct binding or indirect interaction of the PRC2 complex mediated by other proteins such as hnRNPK (Pintacuda et al., 2017). Immunoblotting confirmed a strong and specific interaction of hnRNPK with *Dnmt3bas* (Figure S3B). Overall, these data demonstrate that manipulating *Dnmt3bas* levels significantly affects PRC2 targeting and the deposition of H3K27me3 at the promoter/PE region, that in turn affects the gain of H3K27Ac and *Dnmt3b* induction post-differentiation.

### Dnmt3bas is localized at the Dnmt3b promoter and enhancer and affects E-P looping

We performed a 3C assay in VC, KD, and OE cell lines to test the effect of *Dnmt3bas* on distal enhancer-promoter interaction. The data show little or no effect on the interaction frequency of *Dnmt3b* distal enhancer and promoter in the undifferentiated 2i-ESCs (Figure 4A). However, we observed a higher interaction frequency between enhancer and promoter in KD cells than in VC cells (Figure 4B). These data suggest that an increase in H3K27Ac at *Dnmt3b* regulatory regions reinforces the E-P loop formation in KD cells, potentially affecting the induction level of the *Dnmt3b* gene.

**Figure 4.**
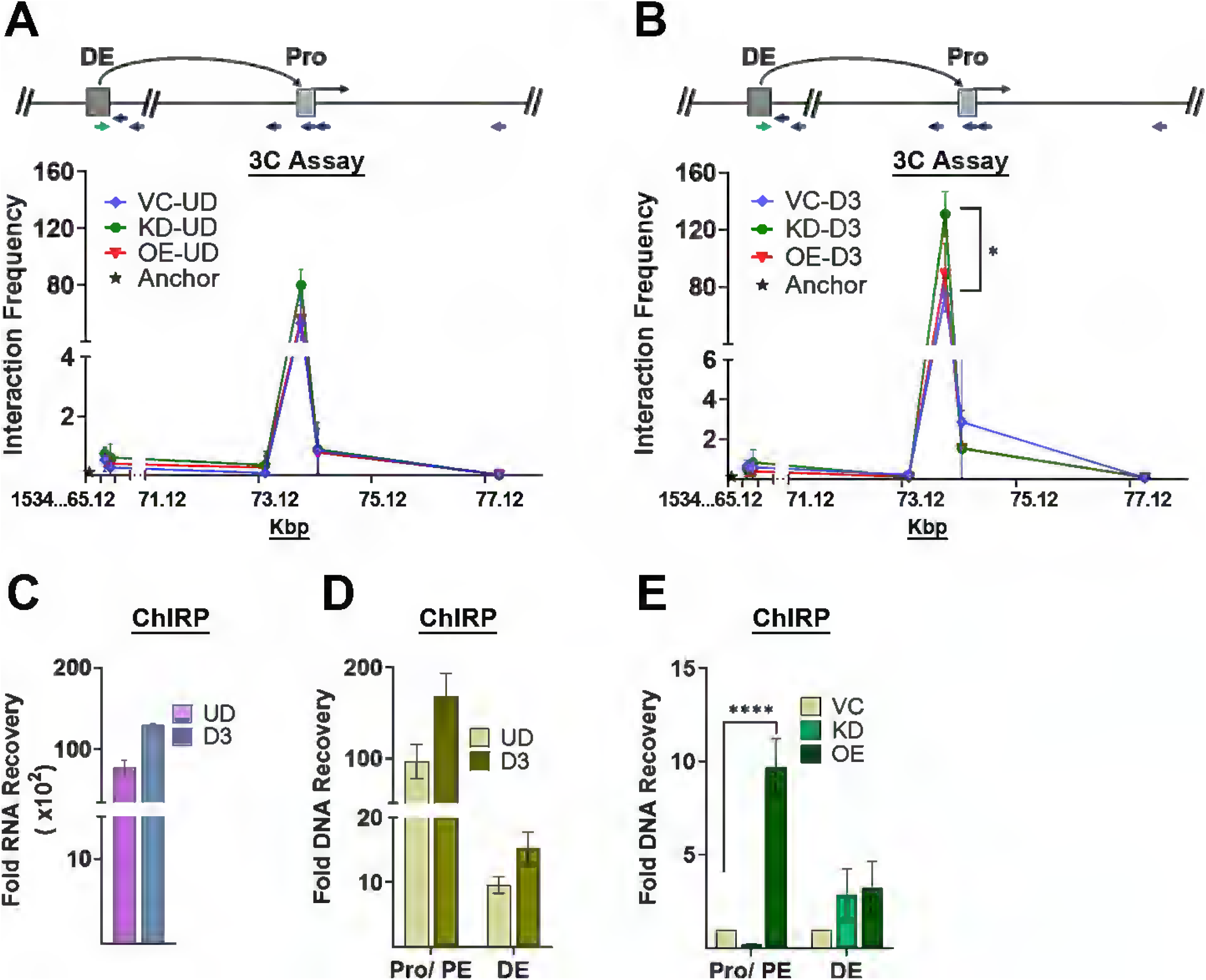
*Dnmt3bas* associates with Dnmt3b enhancers and influences E-P looping. UD: undifferentiated; D3: Days post-induction of differentiation; PE: proximal enhancer; Pro: promoter; DE: Distal enhancer; VC: Vector control; KD: shRNA mediated knockdown of *Dnmt3bas*; OE: overexpression of *Dnmt3bas* lncRNA (A, B) Chromatin interaction assay, 3C, shows an interaction of the distal enhancer with the promoter of Dnmt3b. 3C assays were performed using undifferentiated and D3 differentiated VC, KD, and OE cells. Primer positions were, as described in Figure 1 legend, and primer at DE was used as bait. The data in (A) show no change in E-P interaction in the undifferentiated cells, whereas in (B), a stronger interaction was observed in KD cells at D3 post-differentiation. (C, D, E) ChIRP (chromatin isolation by RNA purification) was performed using *Dnmt3bas*-specific biotinylated probes. The RNA and DNA fractions from the eluate were separated and used to probe for (C) *Dnmt3bas* transcript and (D) Dnmt3b Pro/PE and DE regions, respectively. (C) We observed a prominent recovery of *Dnmt3bas* compared to Gapdh mRNA. (D) From the DNA fraction, the Pro/PE region of Dnmt3b enriched 100-180-fold, and DE showed 10-15-fold recovery compared to a non-specific region in the mouse genome. (E) Fold DNA recovery of Pro/PE and DE regions in *Dnmt3bas* KD and OE cells compared to that in the VC cells. OE cells show a significantly high fold DNA, specifically of the Pro/PE region. Results are presented as normalized mean values ± SEM. P values were derived from Student’s t-test: *p < 0.05; ****p < 0.0001. See also Figure S4.

Given that *Dnmt3bas* affects H3K27Ac at the distal enhancer, we asked if *Dnmt3bas* RNA is associated with the chromatin near the distal enhancer. We performed the Chromatin Isolation by RNA purification (ChIRP) assay in WT and transgenic (VC, OE, and KD) 2iESCs pre- and post-differentiation. Using probes specific to exon or intron regions of *Dnmt3bas*, RNA recovery analysis showed high and specific recovery of the spliced *Dnmt3bas* from the RNA fraction of undifferentiated and D3 differentiated cells (Figure 4C, S4A). From the DNA fraction, approximately 10-fold higher enrichment of the *Dnmt3b* promoter region was measured compared to the distal enhancer (Figure 4D). A significantly lower and higher enrichment of the Dnmt3b promoter region was observed in KD and OE cells, respectively, compared to VC, and no significant change was observed in the enrichment of the Dnmt3b distal enhancer region (Figure 4E). These data confirm that spliced *Dnmt3bas* is located primarily at the *Dnmt3b* promoter.

### *hnRNPL binds Dnmt3bas* at a CA repeat region in exon 2

To determine the mechanism of *Dnmt3bas* in *Dnmt3b* promoter priming and transcriptional induction, we performed an RNA pull-down assay to identify *Dnmt3bas* interacting proteins using i*n vitro* transcribed biotinylated *Dnmt3bas* and control antisense-*Dnmt3bas* RNAs (Figure 5A). The bound proteins were visualized on a Coomassie-stained SDS-PAGE. A distinct protein band at around 60 KDa was explicitly observed in the *Dnmt3bas* elution and was absent in the control sample (Figure S5A). We performed mass spectrometry (MS) analysis of proteins from control and experimental lanes within the excised fragment to identify proteins in the specific band. The data from MS showed a strong enrichment of hnRNPL in the experimental sample. The protein band was confirmed to be hnRNPL by immunoblotting the RNA pull-down samples (Figure 5B). To examine the *in vivo* binding of hnRNPL to *Dnmt3bas*, we performed a western blot analysis of the protein fraction from the *Dnmt3bas* ChIRP assay. The data showed a strong signal of hnRNPL in the experimental sample that was absent in the RNase-treated control sample (Figure 5C). ChIP assay using an anti-hnRNPL antibody showed significant and specific enrichment of hnRNPL at the *Dnmt3b* promoter, confirming its *in vivo* binding at these sites (Figure 5D).

**Figure 5.**
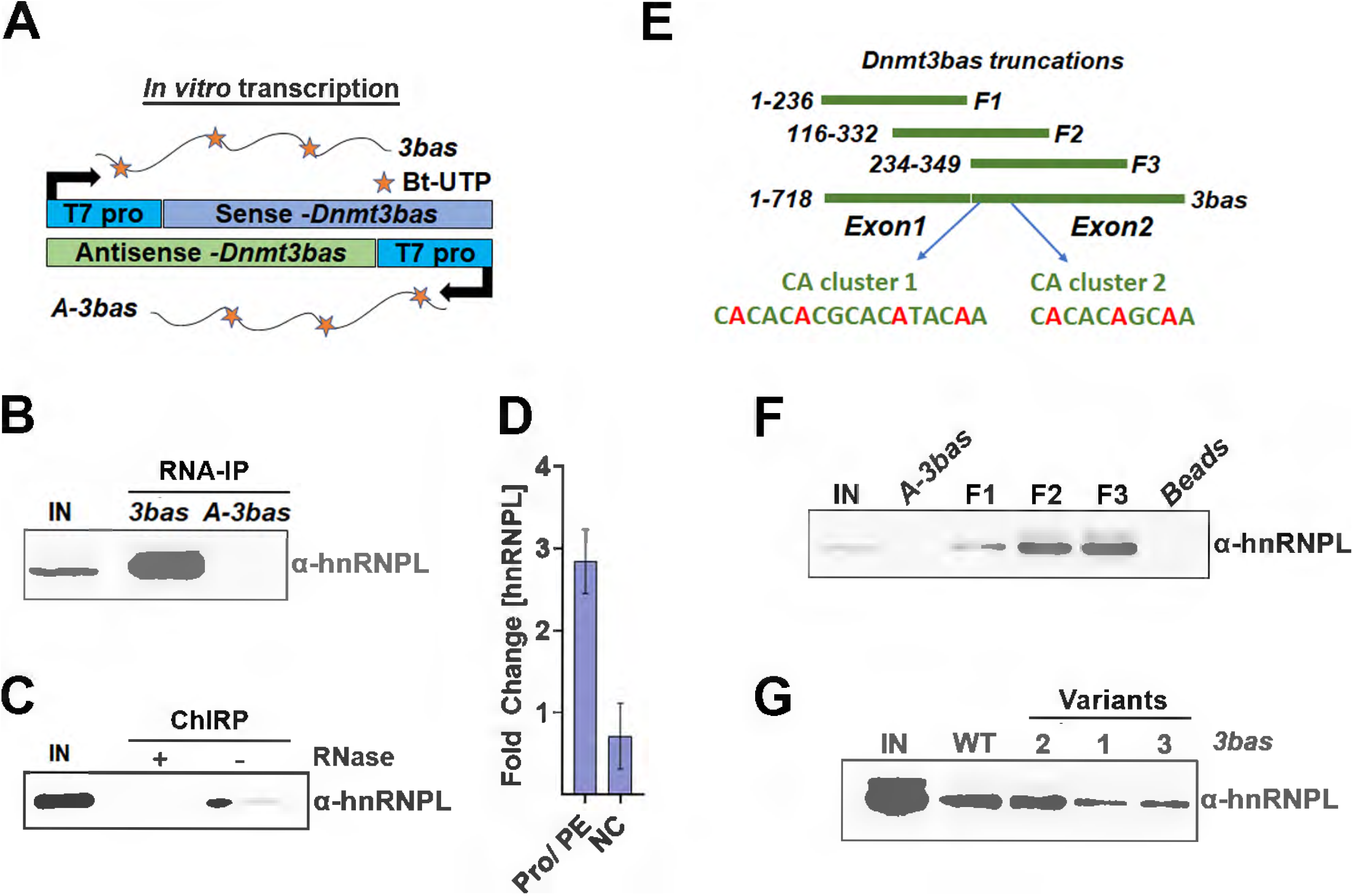
hnRNPL binds to a CA repeat region in exon 2 of *Dnmt3bas*. 3bas: *Dnmt3bas* transcript, A-3bas: Antisense of *Dnmt3bas* transcript, bt-UTP: biotinylated UTP, IN: Input, NC: negative control, E1: Exon1, M: Regions from Exon 1 and Exon 2, E2: Exon2, Cluster 1 and Cluster 2: Predicted hnRNPL binding sites on *Dnmt3bas*, V1, V2: Variant of *Dnmt3bas* transcript with substitutions in cluster 2 (2), and cluster 1 and 2 (1, 3) of hnRNPL binding. (A, E) Schematic showing the experimental design for in vitro transcription of (A) *Dnmt3bas*, antisense control transcript, and (E) various *Dnmt3bas* fragments used for RNA pull-down assays. The *in vitro* transcription was performed using biotinylated UTP and nuclear extract from 2i-ESCs. D) ChIP-qPCR assays show fold enrichment over the input of hnRNPL. NC is the control region in the mouse genome. Results are presented as normalized mean values ± SEM. See also Figure S5 (B, C F, G) Western Blot analysis of hnRNPL; (B) RNA pull-down assay showing specific binding of hnRNPL to *Dnmt3bas* and not to antisense control transcript; (C) Protein fraction from ChIRP assay showing hnRNPL, which was absent in eluate from RNAse treated sample. (F) RNA pull-down assays using fragments of *Dnmt3bas* transcript show strong and specific binding of hnRNPL to a region in Exon 2. (G) Variants of *Dnmt3bas* were used for RNA pull-down assays, showing disrupted interaction of hnRNPL to variants 1 and 3 and not to variant 2

hnRNPL binds CA repeats with high affinity (Smith et al., 2013), and *Dnmt3bas* contains two clusters of CA-repeats within exon 2 spaced apart by about 50bps. To locate the hnRNPL binding sites, we performed RNA pull-down assays using *Dnmt3bas* fragments (Figure 5E). Compared to a weak binding in the exon 1 fragment (F1), strong hnRNPL binding was observed in the intermediate and the exon 2 RNA fragments (F2, F3) containing both CA clusters (Figure 5F). Next, we systematically mutated the CA clusters in the full-length *Dnmt3bas* and used the variants for the RNA pull-down assay. Variants 1 and 3 have different mutations in clusters 1 and 2, and variant 2 has mutations in only cluster 2. Whereas the hnRNPL binding of variant 2 is similar to that of WT *Dnmt3bas*, a very weak binding was detected for variants 1 and 3 (Figure 5G), indicating that hnRNPL predominantly binds to the cluster 1 site. Thus, we conclude that hnRNPL specifically binds *Dnmt3bas* and is recruited to the *Dnmt3b* promoter region.

### *Dnmt3bas* modulates alternative splicing of *Dnmt3b* through hnRNPL activity

Previous studies have shown that the RNA-binding protein hnRNPL regulates alternative splicing (AS) (Cole et al., 2015; Hung et al., 2008; Preussner et al., 2012). Given that *Dnmt3bas* recruits hnRNPL to the *Dnmt3b* promoter, we speculated that post-induction, the *Dnmt3bas*-hnRNPL complex facilitates the interaction of hnRNPL with elongating RNA Pol II, which ferries hnRNPL to alternatively spliced sites at the 3’ end of the *Dnmt3b* transcript. First, we tested the role of *Dnmt3bas* in the alternative splicing of *Dnmt3b. The* ratio of *Dnmt3b1:Dnmt3b6* was determined using RT-PCR analysis in VC, KD, and OE cells cultured in the 2i medium (Figure 6A). *Dnmt3b1:Dnmt3b6* ratio in VC cells pre- and post-differentiation was as shown for WT 2iESCs in Figure 1B. *Dnmt3b6* is the predominant isoform in undifferentiated (UD) 2iESCs. Post differentiation (D3) transcriptional induction of Dnmt3b was accompanied by increased exon inclusion, thus transcribing *Dnmt3b1* as a major isoform. Although this trend was prevalent in KD and OE cells, we observed a peculiar increase in the *Dnmt3b1*: *Dnmt3b6* ratio in OE cells post-differentiation (Figure 6A). To test if increased exon inclusion was due to higher hnRNPL recruitment in OE cells, we overexpressed *Dnmt3bas* variants with mutations in hnRNPL binding sites in 2iESCs. Transgenic cell lines were differentiated, and alternative splicing of *Dnmt3b* was analyzed at D3 post-differentiation. Compared to cells overexpressing WT *Dnmt3bas*, the ratio of *Dnmt3b1*:*Dnmt3b6* in cells overexpressing variants was similar to that in the VC cells (Figure 6B). Furthermore, suppression of hnRNPL expression in OE s-ESCs (serum cultured) rescued the *Dnmt3b6:Dnmt3b1* ratio to that in VC s-ESCs (Figure S6A, 6C). This data confirms the direct role of the hnRNPL/*Dnmt3bas* complex in regulating Dnmt3b alternative splicing.

**Figure 6.**
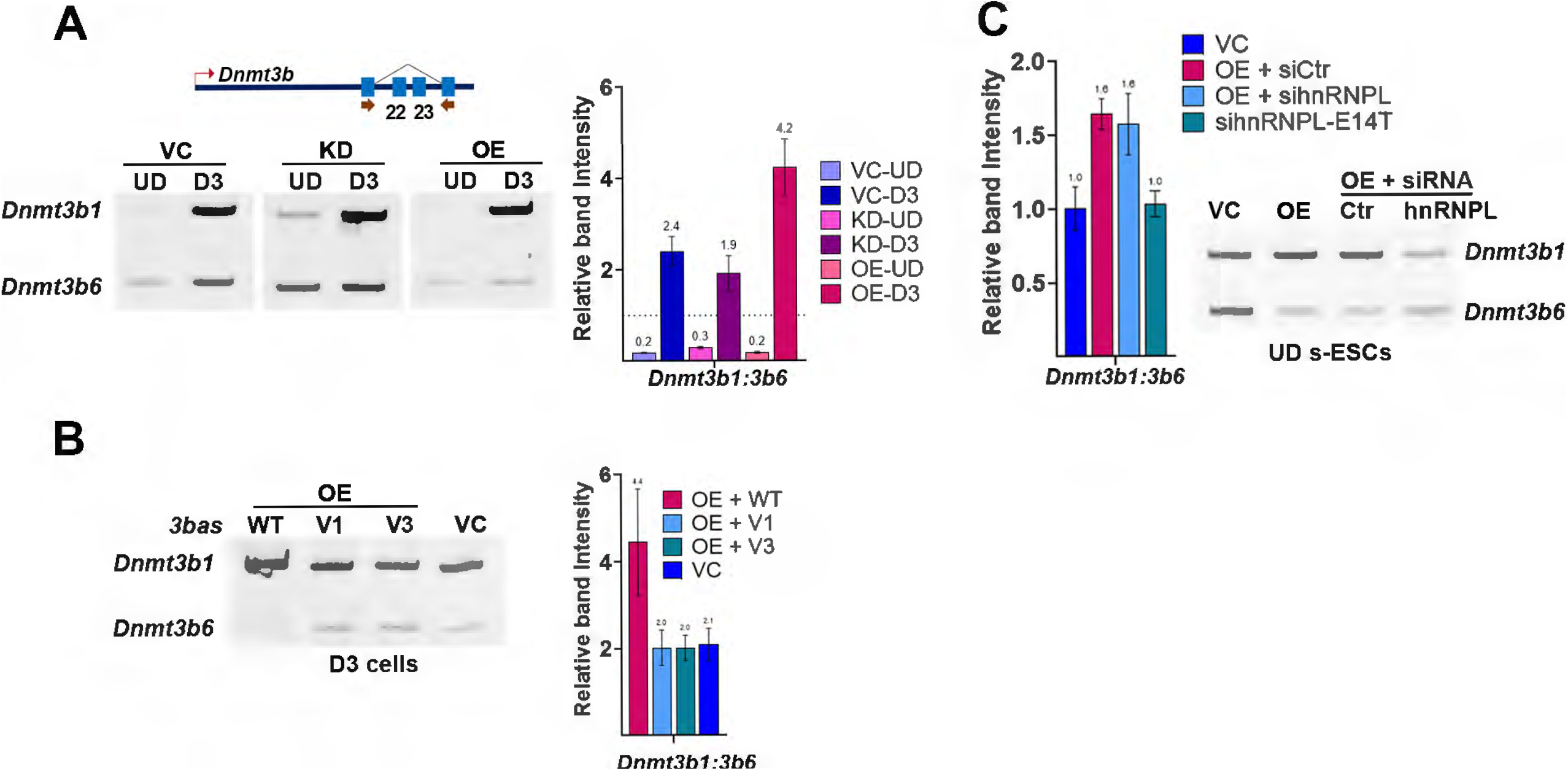
hnRNPL regulates alternative splicing of Dnmt3b. UD: undifferentiated; D3: Days post-induction of differentiation; VC: Vector control; KD: shRNA mediated knockdown of *Dnmt3bas*; OE: overexpression of *Dnmt3bas* lncRNA; V1: Variant 1; V3: variant 3; *Dnmt3b1*: Dnmt3b long isoform (exons 22 and 23 included); *Dnmt3b6*: Dnmt3b short isoform(exons 22 and 23 excluded) s-ESC: serum cultured ESCs (A) RT-PCR analysis of *Dnmt3b1* and *Dnmt3b6* expression amplified in the same reaction by using primers flanking the AS exons. The illustration shows alternatively spliced exons 22 and 23, and red arrows represent the primers’ position to amplify around the alternatively spliced exons of the Dnmt3b gene. The bar graph shows the ratio of quantified band intensity *Dnmt3b1* and *Dnmt3b6*. At D3 post differentiation, a switch to exon inclusion increases *Dnmt3b1* expression relative to *Dnmt3b6*. The ratio of *Dnmt3b1:Dnmt3b6* is further increased in OE cells. B) Ratio of *Dnmt3b1*/*Dnmt3b6* isoforms at D3 from cells overexpressing wildtype *Dnmt3bas*, V1 and V3 mutant *Dnmt3bas*, and vector control. The bar graph shows the ratio of quantified band intensity. (C) siRNA mediated knockdown of hnRNPL in serum cultured ESCs. Control s-ESCs (VC) show an almost equal ratio of *Dnmt3b1*:*Dnmt3b6* isoform expression, whereas Dnmt3bas overexpressing (OE) cells show an enhanced exon inclusion. siRNA-mediated knockdown of hnRNPL in OE cells restored the *Dnmt3b1*:*Dnmt3b6* isoform ratio to 1. Results are presented as normalized mean values ± SEM. See also Figure S6.

Together our data suggest that in *Dnmt3bas* OE cells, increased recruitment of hnRNPL at the promoter increases the number of exon inclusion events. Previous studies have shown that the interaction of SETD2 with hnRNPL and elongating RNA Pol II couples transcriptional elongation and alternative splicing (Bhattacharya et al., 2021a; Bhattacharya et al., 2021b; Yuan et al., 2009). We observed no significant change in the deposition of H3K36me3 at the 3’ end of the *Dnmt3b* gene in *Dnmt3bas* OE cells compared to the VC cells (Figure S6B), confirming that the activities of SETD2 and hnRNPL are not dependent on their interaction with each other (Yuan et al., 2009). However, this observation does not exclude the potential role of SETD2 in mediating the interaction of hnRNPL with elongating Pol II.

### A mechanistic model of Dnmt3bas

Our study comprehensively examined the mechanism that regulates the spatiotemporal expression of *Dnmt3b* during early development (Figure 7). The data revealed the unique role of *Dnmt3bas* in transcriptional priming and coordinated activation of the *Dnmt3b* gene and alternative splicing of the Dnmt*3b* transcript. In the undifferentiated 2i-ESCs, *Dnmt3bas* maintains the *Dnmt3b* promoter in a primed state by regulating PRC2 activity. When differentiation is triggered, *Dnmt3b* expression is induced by enhancement of E-P interaction and deposition of H3K27Ac. Higher expression of *Dnmt3b* is also accompanied by increased exon inclusion and switching of alternative splicing in favor of *Dnmt3b1* transcription. The alternative splicing switch is facilitated by hnRNPL, a *Dnmt3bas* binding protein. At the transcriptionally active *Dnmt3b* promoter, *Dnmt3bas* bridges the interaction of hnRNPL with elongating RNA Pol II, ensuring its delivery to alternative splice sites. Following induction, the post-differentiation repression of *Dnmt3b* is mediated by DNA methylation at proximal and distal enhancers. Notably, the CGI promoter regains the H3K27me3 mark suggesting the role of a lower but persistent expression of *Dnmt3bas*, which is essential for stabilizing the activity of the PRC2 complex at the CGI promoter.

**Figure 7.**
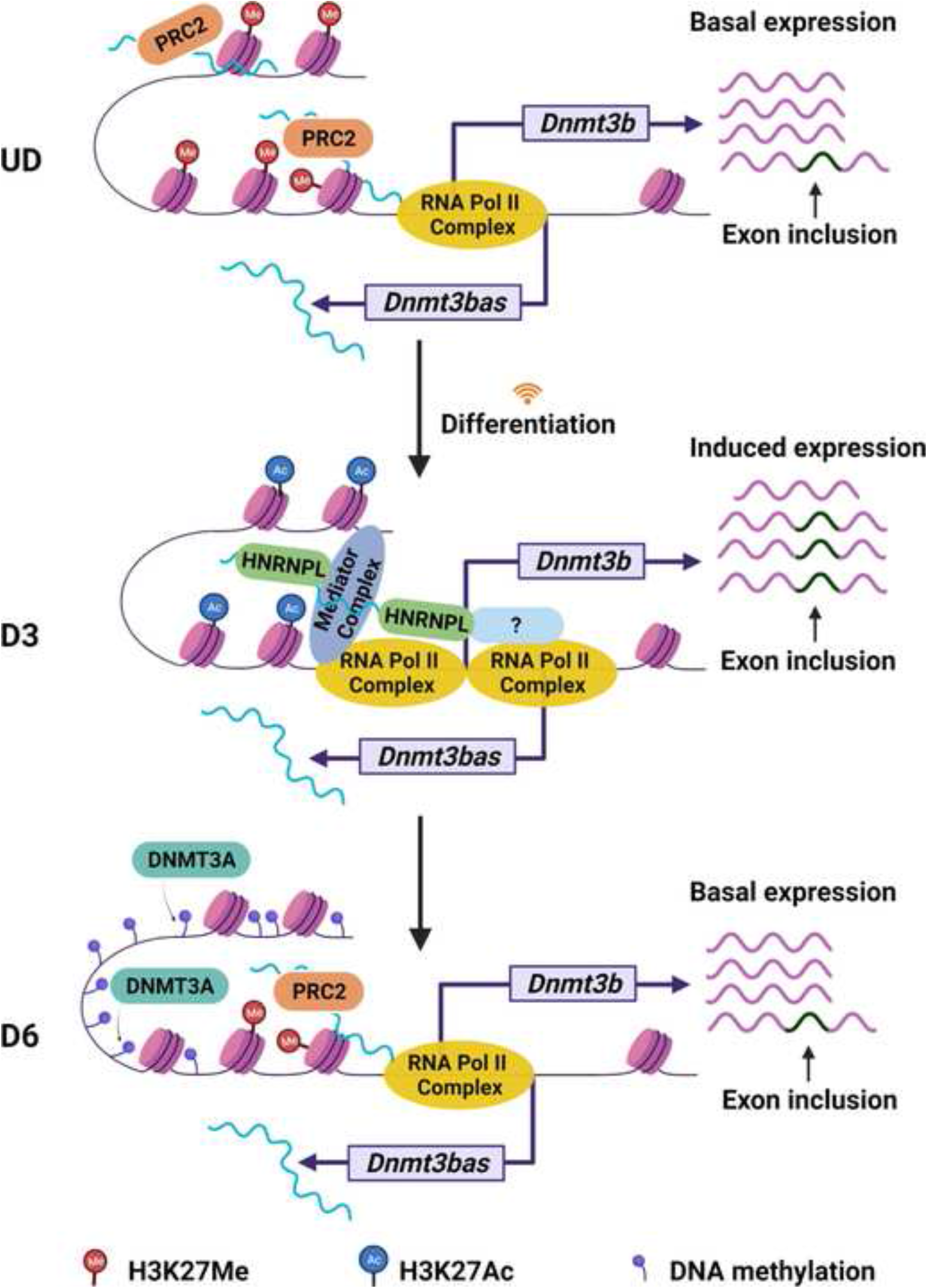
Model of epigenetic changes at pluripotency gene enhancers during stem cell differentiation. In the undifferentiated state (UD), *Dnmt3bas* maintains the *Dntm3b* promoter in a primed state, where the proximal enhancer and promoter regions are characterized by H3K27Me/3 chromatin modifications, facilitating basal *Dnmt3b* expression. In response to differentiation signals (D3), *Dnmt3b* expression is induced via E-P interactions and the deposition of H3K27Ac marks. In concert with induced *Dnmt3b* expression, a switch in alternative splicing increases exon inclusion, promoting *Dnmt3b1* transcription. *Dnmt3bas* directly bind and recruit hnRNPL, an obvious splicing factor, to the transcriptionally active *Dnmt3b* promoter. Once at the promoter, hnRNPL hitchhikes RNA Pol II, potentially through an adaptor protein, ensuring its delivery to *Dnmt3b* splice sites. Post-differentiation (D6), DNA methylation by DNMT3A at the promoter and enhancers reduces the expression of *Dnmt3b* back to basal levels. In particular, the CGI promoter regains H3K27me3 suggesting the role of a lower but persistent expression of *Dnmt3bas*, which is required for stabilizing the activity of the PRC2 complex at the CGI promoter.

## Discussion

Gene expression is controlled by the concerted activity of the transcriptional coactivator complex and a permissive chromatin environment at its cis-regulatory elements. Additionally, divergent long non-coding RNAs (lncRNA) regulate the expression of the nearby protein-coding genes, including those crucial for normal development. They are often expressed from syntenic regions among animal species (Dinger et al., 2008; Ponjavic et al., 2007), and widespread antisense lncRNA transcription is evolutionarily conserved (Seila et al., 2008). LncRNAs act as modulators of gene expression through multiple mechanisms, including the recruitment of chromatin remodelers (Akhade et al.; Guttman et al.). Transcriptional regulation by lncRNAs can be accomplished by either the transcript itself or by the act of transcription, which creates a chromatin environment that influences the expression of the neighboring gene (Gil and Ulitsky, 2020). Many lncRNAs such as HOTAIR (Rinn et al., 2007), Kcnq1ot1 (Thakur et al., 2004), Air (Sleutels et al., 2002), Evx1as(Luo et al., 2016), and Evf2as (Feng et al., 2006) are coordinately transcribed with a protein-coding gene such that the pair is expressed from one transcriptional locus (Engstrom et al., 2006). Several studies have used comparative expression analysis of lncRNA/mRNA pairs to support a functional relationship between the divergent lncRNA and associated protein-coding gene (Sigova et al., 2013). Our study characterized *Dnmt3bas* as a promoter-associated divergent lncRNAs, expressed coordinately with *Dnmt3b* mRNA. Mechanistically, *Dnmt3bas* acts in trans and regulates the induction and alternative splicing of *Dnmt3b*.

### Dnmt3b expression in ESC model system

Serum-cultured ESCs are a mix of two interchangeable cell populations; naïve pluripotent cells and epiblast stem cells. Epiblast stem cells are primed for differentiation, and their gene expression profile shows the expression of early differentiation genes. In contrast, naïve ESCs have high pluripotency genes expression (Tosolini and Jouneau, 2016; Ying et al., 2008). Interestingly, in contrast to serum-ESCs, the expression of *Dnmt3b* is strongly downregulated in 2i-ESCs. Upon induction of differentiation, the transcript levels of *Dnmt3b* increase substantially within a small window of 2-3 days, after which the *Dnmt3b* gene is completely repressed (Ficz et al., 2013; Leitch et al., 2013). This is akin to the dynamic expression of *Dnmt3b* during early embryogenesis that is restricted to the inner mass cells from E4.5-E8.5, after which *Dnmt3b* expression is strongly downregulated in somatic lineages (Hirasawa and Sasaki, 2009; Okano et al., 1999; Watanabe et al., 2002). We used 2i-adapted ESC differentiation to study the dynamic regulation of *Dnmt3b* expression by the combined activity of its cis-regulatory elements and *Dnmt3bas*. During 2i medium adaptation and ESC differentiation, we observed a coordinated yet contrasting pattern of *Dnmt3b* and *Dnmt3bas* expression with the *Dnmt3b* pattern mimicking the expression of *Dnmt3b* observed *in vivo*. The observation that *Dnmt3bas* expression is lowest when the *Dnmt3b* gene is expressed at high levels suggests a potential transcriptional interference mechanism wherein a high rate of sense transcription at the *Dnmt3b* promoter will inhibit the antisense transcription of *Dnmt3bas* (Shuman, 2020). Nevertheless, post-differentiation repression of *Dnmt3b* is not accompanied by an increase in *Dnmt3bas* expression, suggesting a mechanism different from transcriptional interference and confirming the regulatory role of *Dnmt3bas* in maintaining the primed state of the *Dnmt3b* promoter during the pluripotent state.

### Role of Dnmt3bas in Dnmt3b transcriptional induction

Given that *Dnmt3bas* is expressed within the promoter of the *Dnmt3b* gene, genetic manipulations using CRISPR/Cas9 could not be employed since it could have a confounding effect on *Dnmt3b* expression. However, we observed a significant effect of *Dnmt3bas* on *Dnmt3b* expression by using shRNA-mediated knockdown and overexpression of *Dnmt3bas* RNA in the 2i-ESCs. Whereas the overexpression of *Dnmt3bas* suppresses the magnitude of *Dnmt3b* induction, knockdown of *Dnmt3bas* has an opposite effect, complementing the observed inverse correlation between their expression pre-induction and during *Dnmt3b* induction. The chromatin modification state of the *Dnmt3b* cis-regulatory elements shows the impact of *Dnmt3bas* expression on H3K27me3 at CGI promoter and enhancer elements. Our data suggest that *Dnmt3bas* recruits the PRC2 complex to maintain the primed state of the *Dnmt3b* promoter and enhancer elements in undifferentiated 2i-ESCs. In response to the differentiation signal, a concomitant loss of H3K27me3 precedes the gain of H3K27Ac for activation of *Dnmt3b* enhancers, which engage strongly with the promoter to facilitate transcriptional induction.

Some lncRNAs can help stabilize the E-P loop by interacting with components of the Mediator complex like Med1 and Med12 (Luo et al., 2016) or cooperate with CTCF-mediated chromatin interactions to affect the transcription of genes (Saldana-Meyer et al., 2019; Xiang et al., 2014; Yang et al., 2015). However, we observed an increase in the enhancer-promoter (E-P) interaction and H3K27Ac at the distal enhancer in *Dnmt3bas* KD cells, suggesting the role of *Dnmt3bas* in stabilizing the repressive PRC2 complex binding. Furthermore, the proximity of the distal enhancer to the Dnmt3b promoter in undifferentiated cells implies a dual role of the E-P loop, i) in facilitating the recruitment of *Dnmt3bas* and PRC2 complex and ii) mediating a quick transcriptional response to the differentiation signal.

Interestingly neither the expression of *Dnmt3bas* nor enrichment of H3K27me3 at distal regulatory elements increased during the post-differentiation repression of *Dnmt3b*. Instead, we observed a loss of H3K27Ac and a gain of DNA methylation in regions farther upstream, flanking the 5’ end of the CGI promoter. We also showed that DNA methylation was deposited by the methyltransferase Dnmt3a and its allosteric activator protein, Dnmt3l. However, at the CGI promoter, which is maintained in an unmethylated state post-differentiation, we observed an increase in H3K27me3, suggesting the role of the PRC2 complex in maintaining the *Dnmt3b* promoter in a repressed state post-differentiation. This observation also justifies the low but significant expression of *Dnmt3bas* post-differentiation that could be involved in maintaining H3K27me3 at the CGI. Interestingly, DNA methylation at the proximal enhancer region in the OE cells also showed a distinguishable increase post-differentiation, suggesting positive reinforcement of the repressive pathways. These observations are comparable with recent studies showing that the lncRNA potency in spreading the PRC2 complex is tightly linked to lncRNA abundance, genome architecture, and CpG island DNA (Schertzer et al., 2019). We further showed that regulation of Dnmt3b expression by *Dnmt3bas* impacts the gain of DNA methylation globally and at Dnmt3b target sites underscoring the importance of this mechanism.

### Role of Dnmt3bas in Dnmt3b alternative splicing

In ESCs, alternative splicing of the *Dnmt3b* transcript results in two main isoforms: *Dnmt3b6*, exons 22 and 23 are excluded, and *Dnmt3b1*, the full-length transcript (Gowher et al., 2008). Our data here show for the first time that *Dnmt3b* transcripts undergo a splicing switch in 2i-ESCs post-differentiation. Previous studies reporting *Dnmt3b1* as a major isoform used ESCs cultured in FBS. However, the 2i-ESCs, which are a homogenous population of cells in the ground state of pluripotency, express catalytically inactive *Dnmt3b6* as the major isoform, similar to the somatic expression of *Dnmt3b*3. These data suggest that besides downregulated expression, alternative splicing of Dnmt3b contributes to the genome-wide loss of DNA methylation in 2i-ESCs. Concurrently, basal transcription could also retain the Dnmt3b promoter in a primed state for prompt activation. However, this would also necessitate the promoter activation process to append a mechanism that facilitates exon inclusion to generate the full-length *Dnmt3b1*.

Interestingly, among several unique proteins bound to *Dnmt3bas*, RNA pull-down assays identified hnRNPL with the highest specific enrichment. The interaction of hnRNPL with *Dnmt3bas* at the *Dnmt3b* promoter was also confirmed by ChIP and ChIRP assays. HnRNPL functions as a regulator of alternative splicing (Cole et al., 2015; Hung et al., 2008; Preussner et al., 2012), RNA stability (Hui et al., 2003), and transcriptional activation or repression in partnership with multiple lncRNAs (Atianand et al., 2016; Li et al., 2014; Ruan et al., 2016). Previous studies have shown that the binding of hnRNPs, including hnRNPK, hnRNPU, and U1 snRNP binding, facilitate chromatin tethering and nuclear retention of lncRNAs (Hacisuleyman et al., 2016; Lubelsky and Ulitsky, 2018; Shukla et al., 2018; Yin et al., 2020). Med23 was also shown to recruit hnRNPL to the promoter of genes, a role similar to *Dnmt3bas* (Huang et al., 2012). These observations suggest that the nucleoprotein constitution of promoters can determine both transcription and splicing outcomes (Cramer et al., 1997). Functional characterization of *Dnmt3bas*/hnRNPL interaction showed increased inclusion of exons 22 and 23 in *Dnmt3b* transcript in the *Dnmt3bas* OE cells was reversed in cells overexpressing *Dnmt3bas* with a mutation in the hnRNPL binding site. This observation suggests that hnRNPL is the regulator of inducible exon inclusion, a process critical for expressing catalytically active, *Dnmt3b1* during differentiation.

### Implications of Dnmt3bas-hnRNPL interaction

Recent studies have shown the interaction of SETD2 with hnRNPL and proposed that SETD2 hitchhikes to the elongating Pol II and transports hnRNPL to the splice sites as they emerge during chain elongation(Yuan et al., 2009) (Bhattacharya et al., 2021a; Bhattacharya et al., 2021b). Our study suggests that the recruitment of hnRNPL by *Dnmt3bas* to the promoter of *Dnmt3b* facilitates hnRNPL-SETD2 transfer. Given that the RNA binding domain of hnRNPL, RRM2, also interacts with SETD2, the interaction of hnRNPL with SETD2 and *Dnmt3bas* could be mutually exclusive. This prediction is supported by the absence of SETD2 in the RIP/Mass spectrometry analysis of *Dnmt3bas* binding proteins. Therefore, interaction choice could be dictated by the proximity and binding strength of the RRM2 with RNA versus SETD2.

In contrast to hnRNPL, the knockdown of hnRNPI (PTBP1) notably increases exon inclusion in the *Dnmt3b* mRNA (Gowher et al., 2012), suggesting that the antagonistic activity of hnRNPI/hnRNPL potentially determines the splice site choice in *Dnmt3b* pre-mRNA. Both proteins bind to similar polypyrimidine tracts (CA repeats) on RNA through their RRM domains and interact with each other (Hahm et al., 1998). Interestingly PTBP1 is recruited to chromatin by H3K36me3 (Luco et al., 2010), a modification deposited by SETD2. We observed an increase in H3K36me3 post-differentiation (Fig S6B), which could contribute to hnRNPI/hnRNPL recruitment at splice sites. However, the mechanism by which their antagonistic activity at alternative splice sites is regulated is unknown. It would be interesting to determine the mechanism of potential cross-talk between antagonist splice factors and chromatin modifiers that fine-tunes the relative levels of the alternatively spliced isoforms in response to varying physiological conditions. Also, given that *Dnmt3bas* acts in trans, future studies will determine the potential global effect of its activity in regulating expression and alternative splicing of other genes.

## Authors Contributions

M.S.D., I.K.M., M.H., S.M., M.C.H., H.C.W., N.E.B., M.C., M.E., and H.T. performed the experiments. M.C.H., H.G.. analyzed the mass spectrometry data. M.S.D, I.K.M., and H.G. wrote the manuscript

## STAR Methods

### Cell culture, adaptation, differentiation, and transgenic cell lines

As previously described by Alabdi et al., mouse embryonic stem cells (mESCs, E14Tg2A) were cultured and maintained in gelatin-coated tissue culture plates. Differentiation of mESCs was induced by plating 10X10^6^ cells in low attachment 10 cm Petri dishes with a concurrent withdrawal of LIF. After Day 3, embryoid bodies were treated with 1µM Retinoic acid (RA), the medium was replenished every day, cells were harvested each day, and samples were collected for protein, RNA, DNA, and chromatin until Day 9.

CRISPR KO dnmt3a and dnmt3l cell lines were procured from Dr. Taiping Chen, MD Anderson. Transgenic *Dnmt3bas* knockdown (KD) and overexpression (OE) cell lines were generated by transfecting serum-cultured ESCs with pLKO.1 shRNA (Dharmacon) and pcDNA3.1*Dnmt3bas* (spliced and unspliced). pcDNA3.1GFP transfected cells were used as control. After antibiotic selection, the derived stable cell lines were adapted to the 2i culture conditions described below.

### 2i Adaptation

Serum-mESCs were seeded in a T-25 flask and passaged once before adapting them into 2i media. Briefly, the serum-mESCs were washed with PBS and detached with trypsin. FBS was added to inactivate the trypsin, DMEM/F12 media was added, and cells were pelleted. The cells were rewashed to remove any traces of FBS and then plated and grown on gelatin-coated tissue culture plates in 2i media (3μM CHIR99021 and 1μM PD0325901). The cells were adapted until passage 10, and at every passage, the cells were harvested and checked for gene expression and DNA methylation. The differentiation was induced by removing LIF from the media, and at D3, 1µM RA was added, as explained earlier. The samples were harvested at different time points for the analysis.

### Gene expression and alternative splicing analysis by qPCR and RT-PCR

Briefly, total RNA was isolated using the TRIzol reagent (Invitrogen, 15596026) according to the manufacturer’s protocol. First, RNA samples were digested with DNAse (Roche, 04716728001) at 37°C for two hours and purified using a Quick-RNA^TM^ MiniPrep Plus Kit (ZymoReseach, R1057). Then, reverse-transcription quantitative PCR (RT-qPCR) was performed either using Verso One-Step RT-qPCR kits (Thermo Scientific, AB-4104A) or cDNA synthesized using the Tetro cDNA Synthesis Kit (Bioline, BIO-65043). For Verso, we used 5-50ng of RNA per reaction, and to synthesize cDNA, we used 1.2-2.5µg of RNA. cDNA was synthesized either through gene-specific primers (GSP), random hexamers, or oligo (dT) as per the manufacturer’s instructions. Gene expression was calculated as ΔC_t,_ which is C_t_(Gene)-C_t_(*Gapdh or Beta-actin*). Change in gene expression is reported as fold change relative to control undifferentiated cells, which was set to 1, or in a log_2_ scale where values of undifferentiated cells were set to 0. Standard deviations represent at least 2 technical and 2 biological replicates. SEM represents the variance in average data with n=6 or more. The SD, SEM determination, and P-Value were calculated using GraphPad Prism using ANOVA or paired Student T-test. To analyze alternative splicing, a 1 step RT-PCR kit (Invitrogen,) was used to reverse transcribed Trizol purified and DNAse (Roche, 04716728001) treated RNA from ESCs. We designed primers at exon-exon boundaries (View primer list) such that product sizes were 200 – 500 bps long. Supplementary Table S1 details the list of primers used in this study.

### Chromosome Conformation Capture Assay (3C)

3C assays followed the protocol from (Hagege et al., 2007; Naumova et al., 2012) with a few modifications. Murine 2i cells were crosslinked with 1% formaldehyde for 10 minutes at room temperature. Chromatin was suspended in NEB buffer 2.1 and digested with 1000 units HaeIII at 37°C overnight. Crosslinked fragments were ligated with 45 units T4 DNA ligase at 16 °C for 2 hours. A random ligation control template was generated using a BAC clone RP23-474F18 covering the Dnmt3b locus and digested with HaeIII and ligated with T4 ligase. Gapdh locus was used as an endogenous control. Compared to a known concentration of E14 genomic DNA by qPCR with primer pair GaploadF and GaploadR to determine 3C samples concentration. According to the result, adjust each 3C sample to the same template amount for the following quantitative PCR. Interaction frequency between anchor fragment (F1) and distant fragment detection was done in triplicate by quantitative PCR using 3C products and BAC products as a template and 3C primers. Primers position and sequence are indicated in (Fig 1) and Table S1. The relative interaction frequency of each fragment to anchor was calculated as 2^(Ct BAC -Ct 3C)^. P-values were calculated using GraphPad Prism using paired Student T-test.

### RNA-Pulldown and hnRNPL binding assays

The WT and mutant *Dnmt3bas* templates for *in vitro* transcription assays were obtained by PCR using primers containing the T7 promoter sequence (View primer list). Biotin-labeled *Dnmt3bas* transcripts, *Dnmt3bas* antisense, and *Dntm3bas* variants 1, 2, and 3 were transcribed *in vitro* using the T7 RNA polymerase (ThermoFisher Scientific, AM1333) with the following modifications. Unlabeled ATP, GTP, and CTP were used at a concentration of 7.5 mM, except for unlabeled UTP, which was used at 5.63 mM. Notably, the reaction was supplemented with 1.8 mM biotin-UTP (Roche, 11093070910). The transcripts were immobilized on streptavidin beads and incubated with pre-cleared nuclear extract from 2i-ESCs for 45 mins at 4 ^0^C. Following washes, magnetic beads were boiled for 5 mins in the SDS-loading buffer as previously described (Panda et al., 2016). Eluates were analyzed using Coomassie staining and Western blot.

### Subcellular fractionation

20 million mESCs were harvested for the subcellular fractionation, according to (Pandya-Jones and Black, 2009; Wuarin and Schibler, 1994) protocol. Briefly, the cells were trypsinized and centrifuged at low speed at RT for 5min. Cells were lysed by adding the NP-40 lysis buffer for 5 mins. The samples were centrifuged and the supernatant was saved as the cytosolic fraction. The pellet was rinsed with the lysed buffer, to remove any cytoplasmic leftover. The pellet was then resuspended in glycerol buffer by gentle flicking. This was followed by the addition of nuclei lysis buffer. After 2 minutes of incubation, the sample was centrifuged and the supernatant taken as the soluble nuclear fraction/nucleoplasm. Part of the fractions was used to perform the western blot, and the remaining fraction was used to isolate the RNA for RT-qPCR analysis.

### Chromatin Isolation by RNA Purification (ChIRP)

Chromatin Isolation by RNA Purification (ChIRP) was performed by following (Au - Chu et al., 2012). Briefly > 20 million cells were grown and crosslinked 1% glutaraldehyde at room temperature, followed by quenching for 5 minutes. The cell pellets were flash-frozen in liquid nitrogen and stored at -80 °C indefinitely. The cells were thawed at room temperature and dislodged. For 100mg of cell pellet, we used 1ml of lysis buffer containing Protease Inhibitor cocktail, PMSF, and SUPERase•In™ RNase Inhibitor (Invitrogen). The cells were sonicated using a Bioruptor (Diagnode) in 15 ml tubes for 2-3 hours. The sonicated sample was centrifuged, and the supernatant was hybridized with the biotinylated DNA probes specific to lncRNA, *Dnmt3bas*, for 4 hours. In the meantime, C-1 magnetic beads were washed and added to the probe-sample mix. The complex was incubated at 37°C for 30 minutes and then washed 5 times with the 1ml wash buffer. The beads were resuspended in the final wash, and 100ul of the sample was kept for RNA purification. The complex was processed accordingly for the downstream analysis. For RNA purification, the bead complex was resuspended in TriZol, and for DNA isolation, the complex was reverse crosslinked and RNase treated. RNA samples were used to determine fold recovery of *Dnmt3bas* lncRNA by RT-qPCR. The purified ChIRP-DNA was used as a template to determine the enrichment of the genomic region of interest by qPCR. The bead complex was treated with a biotin elution buffer to isolate proteins (Au - Chu et al., 2012) and immunoblotted to identify proteins of interest. Table S2 lists sequences of probes used in this study.

### DNA methylation analysis

*DNA Methylation-Dependent qPCR Assay (MD-qPCR) :* Genomic DNA (gDNA) was purified using the standard phenol-chloroform method. The extracted gDNA was treated with RNAse (Roche) overnight at 37 C. Following another round of Phenol-Chloroform DNA extraction, gDNA was subjected to FspE1 (NEB, R0662S) digestion overnight at 37 C. The digested gDNA was purified by the Phenol-Chloroform method and quantified using the NanoDrop 3300 fluorospectrometer through PicoGreen dye according to the manufacturer’s protocol (Life Technologies, P11495). Quantitative PCR was performed using an equal amount of DNA for each sample. The change in DNA methylation is represented by relative fold change in the Cq value as follows: 2^(Cq(U)-Cq(I)), where Cq(U) is the Cq for the undifferentiated ESC sample, and Cq(I) represents day 3 or day 7 differentiated ESCs. The primers used for DNA methylation-dependent qPCR analysis have been previously described [30]. Standard deviations represent three technical and two biological replicates.

#### DNA Methylation-Dependent Restriction digest (MDR)

Purified genomic DNA was subjected to methylation-sensitive restriction by Hpa II and methylation-insensitive restriction by Msp I overnight at 37 C. Samples were loaded on 0.8% Agarose gel in TAE buffer and bands visualized by Ethidium bromide staining. A smear in the Hpa II digestion lane indicates global loss of DNA methylation.

#### Bisulfite Sequencing

Bisulfite sequencing was performed by using the EpiTect Fast Bisulfite Conversion Kit (Qiagen, 59802) according to the manufacturer’s protocol. Bisulfite-converted DNA was PCR-amplified using published methods (Tremblay et al., 1997) and sequenced using Illumina Wide-SEQ run, which generated over 5K paired-end reads for each sample. The reads were then mapped by Bowtie2 and analyzed by Bismark for DNA methylation. Instances of methylated and unmethylated CpG were quantified and summed to an overall percent methylation for each gene with standard deviations. The significance was calculated by Wilcoxon -matched pairs rank test using GraphPad Prism. See Table S2 for primers used for bisulfite sequencing.

#### Chromatin Immunoprecipitation

ChIP was performed as described (Petell et al., 2016). Briefly, nuclei were isolated from the x-linked cells and were sonicated using Bioruptor (Diagnode), according to the manufacturer’s protocol. A total of 8µg of sheared crosslinked chromatin was incubated with 8µg of antibody pre-loaded on a 1:1 ratio of protein A and protein G magnetic beads (Life Technologies, 10002D and 10004D, respectively). After washing the beads, the samples were eluted in 1% SDS, 10 mM EDTA, 50 mM Tris-HCl, pH 8.0. Crosslinking was reversed by incubation at 65°C for 30 min with shaking. Samples were treated with RNase (Roche, 11119915001) for 2 h at 37°C, and subsequently treated with Proteinase K (Worthington, LS004222) for 2 h at 55°C. DNA was purified by phenol: chloroform extraction followed by ethanol precipitation and quantified using PicoGreen (Life Technologies, P11495) and NanoDrop 3300 fluorospectrometer. qPCR and data analysis were performed as previously described (Petell et al., 2016). Enrichment was calculated as follows = Ct (IN)-Ct (IP) and the fold enrichment over input = 2^ [Ct (IN)-Ct (IP)]. Fold change was calculated by normalizing the fold enrichment at a specific site to that at the control region (Chr 17: 13821873-13821988). The significance of the change was determined via *p*-value, which was calculated by GraphPad Prism using Student’s *t*-test. Supplementary Table S4 lists sequences of primers used.

### Western blot

Standard western blot analysis was conducted on nuclear protein extracts and RNA-pulldown eluates using anti-hnRNP L, 1:1000 (Abcam, ab32680), anti-hnRNP K, 1:1000 (Abclonal, A1701), and anti-rabbit, 1:40,000 (Jackson Immunoresearch, 111-035-003).

Chemiluminescence was performed according to the manufacturer’s protocol (Thermo-Fisher Scientific 34580). For Dnmt3b expression, Protein samples were purified using standard RIPA protein purification (Peach et al., 2015), and blotted with anti-Dnmt3b antibodies (sc-20704), and β-actin (sc-47778) as a loading control. Images were taken using the ChemiDoc MP imaging system (Biorad, 170001402).

## Supplementary figures legends

**Figure S1. Proximal and distal enhancers regulate Dnmt3b induction**

s-ESCs: serum cultured embryonic stem cells; 2i-ESCs: 2i cultured embryonic stem cells 2i: Signaling pathway Inhibitors, CHIR99021, and PD184352; UD: undifferentiated; D3, D6: Days post-induction of differentiation: U: Uncut; H: Hpa II cut; M: Msp I cut

(A, E, F) Relative expression analysis by RT-qPCR of A) various early developmental genes in 2iESCs compared to expression in s-ESCs. E) Relative expression of pluripotency and differentiation genes during 2i-ESC differentiation. The expression of genes in undifferentiated cells was set to 0.

F) Dnmt3b expression in s-ESC pre and post-differentiation. The Ct values were normalized to Gapdh, and expression is shown relative to that in UD cells. The data are an average and standard deviation of three biological replicates.

(B) Alkaline phosphatase staining and SSEA-1 immunofluorescence of s-ESCs and 2i-ESCs. A positive signal indicates pluripotency, which is strong in 2i-ESCs. The scale bar is 100 µm.

(C) DNA methylation analysis. Genomic DNA from 2iESC and s-ESCs was digested with restriction enzymes Hpa II, which cuts unmethylated DNA at the 5’-CCGG-3’ site, and MspI, which cuts DNA at the same site irrespective of the DNA methylation.

(D) Bright field microscopy showing changes in cell morphology as 2i-ESCs differentiate.

G) Western blot showing the expression of Dnmt3b alternative splicing isoforms in undifferentiated (UD) ESCs, Day 3 and Day 6 post-differentiation (D3 and D6, respectively) cells. β-actin was used as the loading control.

H) Schematic of the Dnmt3b gene showing the position of the promoter-associated lncRNA *Dnmt3bas* and upstream enhancer.

F) ChIP-qPCR assays show fold enrichment over the input of H3.

**Figure S2. Spatiotemporal expression of Dnmt3b in *Dnmt3bas* manipulated ESCs**

(A) Illustration of Dnmt3b promoter flanking regions shows the relative position, exon/intron structure, and size of unspliced and spliced transcripts of the lncRNA *Dnmt3bas*, which is transcribed in an antisense direction within the Dnmt3b promoter.

(B) Predicted minimum free energy structure of spliced *Dnmt3bas* transcript, determined using ViennaRNA RNA Fold webserver. The structure is colored by base-pairing probabilities. For unpaired regions, the color denotes the probability of being unpaired, from blue to red going from the highest to the lowest.

C) Relative expression of various genes in 2i-ESCs compared to s-ESCs. The expression of each gene in s-ESCs was set to 1.

D) Ethidium bromide-stained 6% acrylamide gel showing a specific amplification in the qRT-PCR reactions with no signal in the reactions without Reverse Transcriptase (NRT).

E) Relative gene expression of Dnmt3b and *Dnmt3bas* during adaptation (P5-P10) of s-ESCs to 2i medium conditions.

F) Illustration of the transgenic cell line generation and adaptation to 2i media conditions.

G) The bar graph shows an increase in the expression of *Prdm14,* indicating successful adaptation to 2i media conditions.

H) RT-qPCR of *Dnmt3bas* from nuclear and cytosolic RNA fractions from VC, KD and OE cell lines showed about 2 fold higher enrichment in the nucleus compared to the cytosol and a proportional increase in both nuclear and cytoplasmic fractions. Dnmt3b and Gapdh were used as mRNA control, showing higher enrichment in the cytosol than in the nucleus.

I) Brightfield microscopy images of transgenic VC, KD, and OE cells pre and post-differentiation.

J) Relative expression of master pluripotency genes and differentiation-specific genes pre and post-differentiation of the transgenic cell lines, VC, KD, OE.

K) Dnmt3b induction in a transgenic cell line expressing Dnmt3bas shRNA (KD2) during differentiation.

L) DNA methylation analysis. Genomic DNA from transgenic VC, KD and OE cells was digested with restriction enzymes Hpa II, which cuts unmethylated DNA at the 5’-CCGG-3’ site, and MspI, which cuts DNA at the same site irrespective of the DNA methylation. In undifferentiated cells hypomethylated genome is fully digested by Hpa II. However, the post-differentiation gain of DNA methylation protects against digestion by Hpa II prominently in VC cells and less in KD and OE cells.

**Figure S3. Chromatin modification at Dnmt3b regulatory elements in *Dnmt3bas* manipulated ESCs**

A) ChIP-qPCR assay showing fold enrichment of H3K27me3 histone modification at Dnmt3b regulatory elements.

B) Schematic showing the experimental design for in vitro transcription of *Dnmt3bas* and antisense control transcript. The in vitro transcription was performed using biotinylated UTP and nuclear extract from 2i-ESCs. The eluted proteins were analyzed on Western blot to identify hnRNPK and Suz 12.

**Figure S4. *Dnmt3bas* associates with Dnmt3b enhancers and influences E-P looping**

ChIRP was performed using *Dnmt3bas* biotinylated probes specific for spliced and unspliced transcripts. The RNA fraction from the eluate was separated and used to probe for spliced and unspliced *Dnmt3bas* transcript. The data show a 10-fold higher enrichment of spliced transcripts than the unspliced ones.

**Figure S5. hnRNPL binds to a CA repeat region in exon 2 of *Dnmt3bas***

Coomassie-stained gel showing proteins eluted after RNA pull-down using *Dnmt3bas* or anti-*Dnmt3bas* control transcript. The boxes represent the gel slices that were used for the extraction of proteins and detection by mass spectrometry. A prominent band is visible in the *Dnmt3bas* lane compared to the control transcript.

**Figure S6. *Dnmt3bas* associates with Dnmt3b enhancers and influences E-P looping**

(A) Western blot showing the expression of hnRNPL in serum cultured ESC overexpressing *Dnmt3bas* (OE), treated with control siRNA or siRNA to knock down hnRNPL expression. GAPDH was used as the loading control.

(B) ChIP-qPCR assay showing fold enrichment of H3K36me3 histone modifications at the Dnmt3b gene locus.

**Table S1. Primers used in this study.**

A list of all PCR primers used in this study (5’ to 3’), separated by technique

**Table S2. ChIRP probes used in this study.**

A list of all ChIRP probes used in this study (5’ to 3’) with 3’ biotin modification

## Acknowledgments

We are thankful to Gowher lab members for their discussions. This work was supported by NSF award 1716678. In addition, we thank Dr. Tiaping Chen for Dnmt3a^-/-^ and Dnmt3l^-/-^ ESCs. Finally, the authors gratefully acknowledge the Walter Cancer Foundation, DNA Sequencing Facility, and support from the Purdue University Center for Cancer Research, P30CA023168.

## References

1. Akhade, V.S., Pal, D., and Kanduri, C. (2017). Long Noncoding RNA: Genome Organization and Mechanism of Action. Adv Exp Med Biol 1008, 47–74.

2. Atianand, M.K., Hu, W., Satpathy, A.T., Shen, Y., Ricci, E.P., Alvarez-Dominguez, J.R., Bhatta, A., Schattgen, S.A., McGowan, J.D., Blin, J., et al. (2016). A Long Noncoding RNA lincRNA-EPS Acts as a Transcriptional Brake to Restrain Inflammation. Cell 165, 1672–1685.

3. Au - Chu, C., Au - Quinn, J., and Au - Chang, H.Y. (2012). Chromatin Isolation by RNA Purification (ChIRP). JoVE, e3912.

4. Bhattacharya, S., Levy, M.J., Zhang, N., Li, H., Florens, L., Washburn, M.P., and Workman, J.L. (2021a). The methyltransferase SETD2 couples transcription and splicing by engaging mRNA processing factors through its SHI domain. Nat Commun 12, 1443.

5. Bhattacharya, S., Wang, S., Reddy, D., Shen, S., Zhang, Y., Zhang, N., Li, H., Washburn, M.P., Florens, L., Shi, Y., et al. (2021b). Structural basis of the interaction between SETD2 methyltransferase and hnRNP L paralogs for governing co-transcriptional splicing. Nat Commun 12, 6452.

6. Cole, B.S., Tapescu, I., Allon, S.J., Mallory, M.J., Qiu, J., Lake, R.J., Fan, H.Y., Fu, X.D., and Lynch, K.W. (2015). Global analysis of physical and functional RNA targets of hnRNP L reveals distinct sequence and epigenetic features of repressed and enhanced exons. RNA 21, 2053–2066.

7. Cramer, P., Pesce, C.G., Baralle, F.E., and Kornblihtt, A.R. (1997). Functional association between promoter structure and transcript alternative splicing. Proc Natl Acad Sci U S A 94, 11456–11460.

8. Derrien, T., Johnson, R., Bussotti, G., Tanzer, A., Djebali, S., Tilgner, H., Guernec, G., Martin, D., Merkel, A., Knowles, D.G., et al. (2012). The GENCODE v7 catalog of human long noncoding RNAs: analysis of their gene structure, evolution, and expression. Genome Res 22, 1775–1789.

9. Dinger, M.E., Pang, K.C., Mercer, T.R., and Mattick, J.S. (2008). Differentiating protein-coding and noncoding RNA: challenges and ambiguities. PLoS computational biology 4, e1000176.

10. Duymich, C.E., Charlet, J., Yang, X., Jones, P.A., and Liang, G. (2016). DNMT3B isoforms without catalytic activity stimulate gene body methylation as accessory proteins in somatic cells. Nat Commun 7, 11453.

11. Engstrom, P.G., Suzuki, H., Ninomiya, N., Akalin, A., Sessa, L., Lavorgna, G., Brozzi, A., Luzi, L., Tan, S.L., Yang, L., et al. (2006). Complex Loci in human and mouse genomes. PLoS genetics 2, e47.

12. Esteller, M. (2005). Aberrant DNA methylation as a cancer-inducing mechanism. Annual review of pharmacology and toxicology 45, 629–656.

13. Feng, J., Bi, C., Clark, B.S., Mady, R., Shah, P., and Kohtz, J.D. (2006). The Evf-2 noncoding RNA is transcribed from the Dlx-5/6 ultraconserved region and functions as a Dlx-2 transcriptional coactivator. Genes Dev 20, 1470–1484.

14. Ficz, G., Hore, T.A., Santos, F., Lee, H.J., Dean, W., Arand, J., Krueger, F., Oxley, D., Paul, Y.L., Walter, J., et al. (2013). FGF signaling inhibition in ESCs drives rapid genome-wide demethylation to the epigenetic ground state of pluripotency. Cell Stem Cell 13, 351–359.

15. Gil, N., and Ulitsky, I. (2020). Regulation of gene expression by cis-acting long non-coding RNAs. Nat Rev Genet 21, 102–117.

16. Gopalakrishnan, S., Sullivan, B.A., Trazzi, S., Della Valle, G., and Robertson, K.D. (2009). DNMT3B interacts with constitutive centromere protein CENP-C to modulate DNA methylation and the histone code at centromeric regions. Human molecular genetics 18, 3178–3193.

17. Gowher, H., Brick, K., Camerini-Otero, R.D., and Felsenfeld, G. (2012). Vezf1 protein binding sites genome-wide are associated with pausing of elongating RNA polymerase II. Proc Natl Acad Sci U S A 109, 2370–2375.

18. Gowher, H., and Jeltsch, A. (2002). Molecular enzymology of the catalytic domains of the Dnmt3a and Dnmt3b DNA methyltransferases. The Journal of biological chemistry 277, 20409–20414.

19. Gowher, H., Stuhlmann, H., and Felsenfeld, G. (2008). Vezf1 regulates genomic DNA methylation through its effects on expression of DNA methyltransferase Dnmt3b. Genes Dev 22, 2075–2084.

20. Greenberg, M.V.C., and Bourc’his, D. (2019). The diverse roles of DNA methylation in mammalian development and disease. Nature Reviews Molecular Cell Biology 20, 590–607.

21. Guil, S., and Esteller, M. (2009). DNA methylomes, histone codes and miRNAs: tying it all together. The international journal of biochemistry & cell biology 41, 87–95.

22. Guo, C.J., Ma, X.K., Xing, Y.H., Zheng, C.C., Xu, Y.F., Shan, L., Zhang, J., Wang, S., Wang, Y., Carmichael, G.G., et al. (2020). Distinct Processing of lncRNAs Contributes to Non-conserved Functions in Stem Cells. Cell 181, 621–636 e622.

23. Guttman, M., Amit, I., Garber, M., French, C., Lin, M.F., Feldser, D., Huarte, M., Zuk, O., Carey, B.W., Cassady, J.P., et al. (2009). Chromatin signature reveals over a thousand highly conserved large non- coding RNAs in mammals. Nature 458, 223–227.

24. Guttman, M., Donaghey, J., Carey, B.W., Garber, M., Grenier, J.K., Munson, G., Young, G., Lucas, A.B., Ach, R., Bruhn, L., et al. (2011). lincRNAs act in the circuitry controlling pluripotency and differentiation. Nature 477, 295–300.

25. Hacisuleyman, E., Shukla, C.J., Weiner, C.L., and Rinn, J.L. (2016). Function and evolution of local repeats in the Firre locus. Nat Commun 7, 11021.

26. Hagege, H., Klous, P., Braem, C., Splinter, E., Dekker, J., Cathala, G., de Laat, W., and Forne, T. (2007). Quantitative analysis of chromosome conformation capture assays (3C-qPCR). Nat Protoc 2, 1722–1733.

27. Hahm, B., Cho, O.H., Kim, J.E., Kim, Y.K., Kim, J.H., Oh, Y.L., and Jang, S.K. (1998). Polypyrimidine tract-binding protein interacts with HnRNP L. FEBS Lett 425, 401–406.

28. Hirasawa, R., and Sasaki, H. (2009). Dynamic transition of Dnmt3b expression in mouse pre- and early post-implantation embryos. Gene Expr Patterns 9, 27–30.

29. Huang, Y., Li, W., Yao, X., Lin, Q.J., Yin, J.W., Liang, Y., Heiner, M., Tian, B., Hui, J., and Wang, G. (2012). Mediator complex regulates alternative mRNA processing via the MED23 subunit. Mol Cell 45, 459–469.

30. Hui, J., Reither, G., and Bindereif, A. (2003). Novel functional role of CA repeats and hnRNP L in RNA stability. RNA 9, 931–936.

31. Hung, L.H., Heiner, M., Hui, J., Schreiner, S., Benes, V., and Bindereif, A. (2008). Diverse roles of hnRNP L in mammalian mRNA processing: a combined microarray and RNAi analysis. RNA 14, 284–296.

32. Huntriss, J., Hinkins, M., Oliver, B., Harris, S.E., Beazley, J.C., Rutherford, A.J., Gosden, R.G., Lanzendorf, S.E., and Picton, H.M. (2004). Expression of mRNAs for DNA methyltransferases and methyl-CpG-binding proteins in the human female germ line, preimplantation embryos, and embryonic stem cells. Molecular reproduction and development 67, 323–336.

33. Ishida, C., Ura, K., Hirao, A., Sasaki, H., Toyoda, A., Sakaki, Y., Niwa, H., Li, E., and Kaneda, Y. (2003). Genomic organization and promoter analysis of the Dnmt3b gene. Gene 310, 151–159.

34. Jinawath, A., Miyake, S., Yanagisawa, Y., Akiyama, Y., and Yuasa, Y. (2005). Transcriptional regulation of the human DNA methyltransferase 3A and 3B genes by Sp3 and Sp1 zinc finger proteins. The Biochemical journal 385, 557–564.

35. Kulis, M., and Esteller, M. (2010). DNA methylation and cancer. Advances in genetics 70, 27–56.

36. Leitch, H.G., McEwen, K.R., Turp, A., Encheva, V., Carroll, T., Grabole, N., Mansfield, W., Nashun, B., Knezovich, J.G., Smith, A., et al. (2013). Naive pluripotency is associated with global DNA hypomethylation. Nature structural & molecular biology 20, 311–316.

37. Li, Z., Chao, T.C., Chang, K.Y., Lin, N., Patil, V.S., Shimizu, C., Head, S.R., Burns, J.C., and Rana, T.M. (2014). The long noncoding RNA THRIL regulates TNFalpha expression through its interaction with hnRNPL. Proc Natl Acad Sci U S A 111, 1002–1007.

38. Lubelsky, Y., and Ulitsky, I. (2018). Sequences enriched in Alu repeats drive nuclear localization of long RNAs in human cells. Nature 555, 107–111.

39. Luco, R.F., Pan, Q., Tominaga, K., Blencowe, B.J., Pereira-Smith, O.M., and Misteli, T. (2010). Regulation of alternative splicing by histone modifications. Science 327, 996–1000.

40. Luo, S., Lu, J.Y., Liu, L., Yin, Y., Chen, C., Han, X., Wu, B., Xu, R., Liu, W., Yan, P., et al. (2016). Divergent lncRNAs Regulate Gene Expression and Lineage Differentiation in Pluripotent Cells. Cell Stem Cell 18, 637–652.

41. Mele, M., Mattioli, K., Mallard, W., Shechner, D.M., Gerhardinger, C., and Rinn, J.L. (2017). Chromatin environment, transcriptional regulation, and splicing distinguish lincRNAs and mRNAs. Genome Res 27, 27–37.

42. Mensah, I.K., Norvil, A.B., AlAbdi, L., McGovern, S., Petell, C.J., He, M., and Gowher, H. (2021). Misregulation of the expression and activity of DNA methyltransferases in cancer. NAR Cancer 3, zcab045.

43. Montgomery, K.G., Liu, M.C., Eccles, D.M., and Campbell, I.G. (2004). The DNMT3B C-->T promoter polymorphism and risk of breast cancer in a British population: a case-control study. Breast cancer research : BCR 6, R390–394.

44. Naumova, N., Smith, E.M., Zhan, Y., and Dekker, J. (2012). Analysis of long-range chromatin interactions using Chromosome Conformation Capture. Methods 58, 192–203.

45. Norvil, A.B., AlAbdi, L., Liu, B., Tu, Y.H., Forstoffer, N.E., Michie, A.R., Chen, T., and Gowher, H. (2020). The acute myeloid leukemia variant DNMT3A Arg882His is a DNMT3B-like enzyme. Nucleic Acids Res 48, 3761–3775.

46. Okano, M., Bell, D.W., Haber, D.A., and Li, E. (1999). DNA methyltransferases Dnmt3a and Dnmt3b are essential for de novo methylation and mammalian development. Cell 99, 247–257.

47. Okano, M., Xie, S., and Li, E. (1998). Cloning and characterization of a family of novel mammalian DNA (cytosine-5) methyltransferases. Nature genetics 19, 219–220.

48. Ostler, K.R., Davis, E.M., Payne, S.L., Gosalia, B.B., Exposito-Cespedes, J., Le Beau, M.M., and Godley, L.A. (2007). Cancer cells express aberrant DNMT3B transcripts encoding truncated proteins. Oncogene 26, 5553–5563.

49. Panda, A.C., Martindale, J.L., and Gorospe, M. (2016). Affinity Pulldown of Biotinylated RNA for Detection of Protein-RNA Complexes. Bio-protocol 6, e2062.

50. Pandya-Jones, A., and Black, D.L. (2009). Co-transcriptional splicing of constitutive and alternative exons. Rna 15, 1896–1908.

51. Peach, M., Marsh, N., Miskiewicz, E.I., and MacPhee, D.J. (2015). Solubilization of proteins: the importance of lysis buffer choice. Methods Mol Biol 1312, 49–60.

52. Petell, C.J., Alabdi, L., He, M., San Miguel, P., Rose, R., and Gowher, H. (2016). An epigenetic switch regulates de novo DNA methylation at a subset of pluripotency gene enhancers during embryonic stem cell differentiation. Nucleic Acids Res 44, 7605–7617.

53. Pintacuda, G., Wei, G., Roustan, C., Kirmizitas, B.A., Solcan, N., Cerase, A., Castello, A., Mohammed, S., Moindrot, B., Nesterova, T.B., et al. (2017). hnRNPK Recruits PCGF3/5-PRC1 to the Xist RNA B-Repeat to Establish Polycomb-Mediated Chromosomal Silencing. Mol Cell 68, 955–969.e910.

54. Ponjavic, J., Ponting, C.P., and Lunter, G. (2007). Functionality or transcriptional noise? Evidence for selection within long noncoding RNAs. Genome Res 17, 556–565.

55. Preussner, M., Schreiner, S., Hung, L.H., Porstner, M., Jack, H.M., Benes, V., Ratsch, G., and Bindereif, A. (2012). HnRNP L and L-like cooperate in multiple-exon regulation of CD45 alternative splicing. Nucleic acids research 40, 5666–5678.

56. Rinn, J.L., Kertesz, M., Wang, J.K., Squazzo, S.L., Xu, X., Brugmann, S.A., Goodnough, L.H., Helms, J.A., Farnham, P.J., Segal, E., et al. (2007). Functional demarcation of active and silent chromatin domains in human HOX loci by noncoding RNAs. Cell 129, 1311–1323.

57. Robertson, K.D., Keyomarsi, K., Gonzales, F.A., Velicescu, M., and Jones, P.A. (2000). Differential mRNA expression of the human DNA methyltransferases (DNMTs) 1, 3a and 3b during the G(0)/G(1) to S phase transition in normal and tumor cells. Nucleic acids research 28, 2108–2113.

58. Robertson, K.D., Uzvolgyi, E., Liang, G., Talmadge, C., Sumegi, J., Gonzales, F.A., and Jones, P.A. (1999). The human DNA methyltransferases (DNMTs) 1, 3a and 3b: coordinate mRNA expression in normal tissues and overexpression in tumors. Nucleic acids research 27, 2291–2298.

59. Ruan, X., Li, P., Cangelosi, A., Yang, L., and Cao, H. (2016). A Long Non-coding RNA, lncLGR, Regulates Hepatic Glucokinase Expression and Glycogen Storage during Fasting. Cell Rep 14, 1867–1875.

60. Saito, Y., Kanai, Y., Sakamoto, M., Saito, H., Ishii, H., and Hirohashi, S. (2002). Overexpression of a splice variant of DNA methyltransferase 3b, DNMT3b4, associated with DNA hypomethylation on pericentromeric satellite regions during human hepatocarcinogenesis. Proc Natl Acad Sci U S A 99, 10060–10065.

61. Saldana-Meyer, R., Rodriguez-Hernaez, J., Escobar, T., Nishana, M., Jacome-Lopez, K., Nora, E.P., Bruneau, B.G., Tsirigos, A., Furlan-Magaril, M., Skok, J., et al. (2019). RNA Interactions Are Essential for CTCF-Mediated Genome Organization. Mol Cell 76, 412–422 e415.

62. Schertzer, M.D., Braceros, K.C.A., Starmer, J., Cherney, R.E., Lee, D.M., Salazar, G., Justice, M., Bischoff, S.R., Cowley, D.O., Ariel, P., et al. (2019). lncRNA-Induced Spread of Polycomb Controlled by Genome Architecture, RNA Abundance, and CpG Island DNA. Mol Cell 75, 523–537 e510.

63. Seila, A.C., Calabrese, J.M., Levine, S.S., Yeo, G.W., Rahl, P.B., Flynn, R.A., Young, R.A., and Sharp, P.A. (2008). Divergent transcription from active promoters. Science 322, 1849–1851.

64. Shen, H., Wang, L., Spitz, M.R., Hong, W.K., Mao, L., and Wei, Q. (2002). A novel polymorphism in human cytosine DNA-methyltransferase-3B promoter is associated with an increased risk of lung cancer. Cancer research 62, 4992–4995.

65. Shukla, C.J., McCorkindale, A.L., Gerhardinger, C., Korthauer, K.D., Cabili, M.N., Shechner, D.M., Irizarry, R.A., Maass, P.G., and Rinn, J.L. (2018). High-throughput identification of RNA nuclear enrichment sequences. EMBO J 37.

66. Shuman, S. (2020). Transcriptional interference at tandem lncRNA and protein-coding genes: an emerging theme in regulation of cellular nutrient homeostasis. Nucleic acids research 48, 8243–8254.

67. Sigova, A.A., Mullen, A.C., Molinie, B., Gupta, S., Orlando, D.A., Guenther, M.G., Almada, A.E., Lin, C., Sharp, P.A., Giallourakis, C.C., et al. (2013). Divergent transcription of long noncoding RNA/mRNA gene pairs in embryonic stem cells. Proc Natl Acad Sci U S A 110, 2876–2881.

68. Singal, R., Das, P.M., Manoharan, M., Reis, I.M., and Schlesselman, J.J. (2005). Polymorphisms in the DNA methyltransferase 3b gene and prostate cancer risk. Oncology reports 14, 569–573.

69. Sleutels, F., Zwart, R., and Barlow, D.P. (2002). The non-coding Air RNA is required for silencing autosomal imprinted genes. Nature 415, 810–813.

70. Smith, S.A., Ray, D., Cook, K.B., Mallory, M.J., Hughes, T.R., and Lynch, K.W. (2013). Paralogs hnRNP L and hnRNP LL exhibit overlapping but distinct RNA binding constraints. PLoS One 8, e80701.

71. Statello, L., Guo, C.J., Chen, L.L., and Huarte, M. (2021). Gene regulation by long non-coding RNAs and its biological functions. Nat Rev Mol Cell Biol 22, 96–118.

72. Szyf, M. (2005). Therapeutic implications of DNA methylation. Future Oncol 1, 125–135.

73. Thakur, N., Tiwari, V.K., Thomassin, H., Pandey, R.R., Kanduri, M., Gondor, A., Grange, T., Ohlsson, R., and Kanduri, C. (2004). An antisense RNA regulates the bidirectional silencing property of the Kcnq1 imprinting control region. Molecular and cellular biology 24, 7855–7862.

74. Tian, B., and Manley, J.L. (2017). Alternative polyadenylation of mRNA precursors. Nat Rev Mol Cell Biol 18, 18–30.

75. Tosolini, M., and Jouneau, A. (2016). Acquiring Ground State Pluripotency: Switching Mouse Embryonic Stem Cells from Serum/LIF Medium to 2i/LIF Medium. Methods Mol Biol 1341, 41–48.

76. Tremblay, K.D., Duran, K.L., and Bartolomei, M.S. (1997). A 5’ 2-kilobase-pair region of the imprinted mouse H19 gene exhibits exclusive paternal methylation throughout development. Molecular and cellular biology 17, 4322–4329.

77. Walton, E.L., Francastel, C., and Velasco, G. (2014). Dnmt3b Prefers Germ Line Genes and Centromeric Regions: Lessons from the ICF Syndrome and Cancer and Implications for Diseases. Biology 3, 578–605.

78. Wang, J., Bhutani, M., Pathak, A.K., Lang, W., Ren, H., Jelinek, J., He, R., Shen, L., Issa, J.P., and Mao, L. (2007). Delta DNMT3B variants regulate DNA methylation in a promoter-specific manner. Cancer research 67, 10647–10652.

79. Watanabe, D., Suetake, I., Tada, T., and Tajima, S. (2002). Stage- and cell-specific expression of Dnmt3a and Dnmt3b during embryogenesis. Mech Dev 118, 187–190.

80. Watanabe, Y., and Maekawa, M. (2010). Methylation of DNA in cancer. Advances in clinical chemistry 52, 145–167.

81. Weisenberger, D.J., Velicescu, M., Cheng, J.C., Gonzales, F.A., Liang, G., and Jones, P.A. (2004). Role of the DNA methyltransferase variant DNMT3b3 in DNA methylation. Mol Cancer Res 2, 62–72.

82. Wuarin, J., and Schibler, U. (1994). Physical isolation of nascent RNA chains transcribed by RNA polymerase II: evidence for cotranscriptional splicing. Mol Cell Biol 14, 7219–7225.

83. Xiang, J.F., Yin, Q.F., Chen, T., Zhang, Y., Zhang, X.O., Wu, Z., Zhang, S., Wang, H.B., Ge, J., Lu, X., et al. (2014). Human colorectal cancer-specific CCAT1-L lncRNA regulates long-range chromatin interactions at the MYC locus. Cell Res 24, 513–531.

84. Xie, S., Wang, Z., Okano, M., Nogami, M., Li, Y., He, W.W., Okumura, K., and Li, E. (1999). Cloning, expression and chromosome locations of the human DNMT3 gene family. Gene 236, 87–95.

85. Xie, Z.H., Huang, Y.N., Chen, Z.X., Riggs, A.D., Ding, J.P., Gowher, H., Jeltsch, A., Sasaki, H., Hata, K., and Xu, G.L. (2006). Mutations in DNA methyltransferase DNMT3B in ICF syndrome affect its regulation by DNMT3L. Hum Mol Genet 15, 1375–1385.

86. Xu, G.L., Bestor, T.H., Bourc’his, D., Hsieh, C.L., Tommerup, N., Bugge, M., Hulten, M., Qu, X., Russo, J.J., and Viegas-Pequignot, E. (1999). Chromosome instability and immunodeficiency syndrome caused by mutations in a DNA methyltransferase gene. Nature 402, 187–191.

87. Yagi, M., Kabata, M., Tanaka, A., Ukai, T., Ohta, S., Nakabayashi, K., Shimizu, M., Hata, K., Meissner, A., Yamamoto, T., et al. (2020). Identification of distinct loci for de novo DNA methylation by DNMT3A and DNMT3B during mammalian development. Nature Communications 11, 3199.

88. Yang, F., Deng, X., Ma, W., Berletch, J.B., Rabaia, N., Wei, G., Moore, J.M., Filippova, G.N., Xu, J., Liu, Y., et al. (2015). The lncRNA Firre anchors the inactive X chromosome to the nucleolus by binding CTCF and maintains H3K27me3 methylation. Genome biology 16, 52.

89. Yin, Y., Lu, J.Y., Zhang, X., Shao, W., Xu, Y., Li, P., Hong, Y., Cui, L., Shan, G., Tian, B., et al. (2020). U1 snRNP regulates chromatin retention of noncoding RNAs. Nature 580, 147–150.

90. Ying, Q.L., Wray, J., Nichols, J., Batlle-Morera, L., Doble, B., Woodgett, J., Cohen, P., and Smith, A. (2008). The ground state of embryonic stem cell self-renewal. Nature 453, 519–523.

91. Yuan, W., Xie, J., Long, C., Erdjument-Bromage, H., Ding, X., Zheng, Y., Tempst, P., Chen, S., Zhu, B., and Reinberg, D. (2009). Heterogeneous nuclear ribonucleoprotein L Is a subunit of human KMT3a/Set2 complex required for H3 Lys-36 trimethylation activity in vivo. The Journal of biological chemistry 284, 15701–15707.

92. Zeng, Y., Ren, R., Kaur, G., Hardikar, S., Ying, Z., Babcock, L., Gupta, E., Zhang, X., Chen, T., and Cheng, X. (2020). The inactive Dnmt3b3 isoform preferentially enhances Dnmt3b-mediated DNA methylation. Genes Dev 34, 1546–1558.

93. Zuckerman, B., and Ulitsky, I. (2019). Predictive models of subcellular localization of long RNAs. RNA 25, 557–572.

