## Supplementary figures and text for "*Dnmt3bas* coordinates transcriptional induction and alternative exon inclusion to promote catalytically active Dnmt3b expression"

#### **Supplemental Figures**

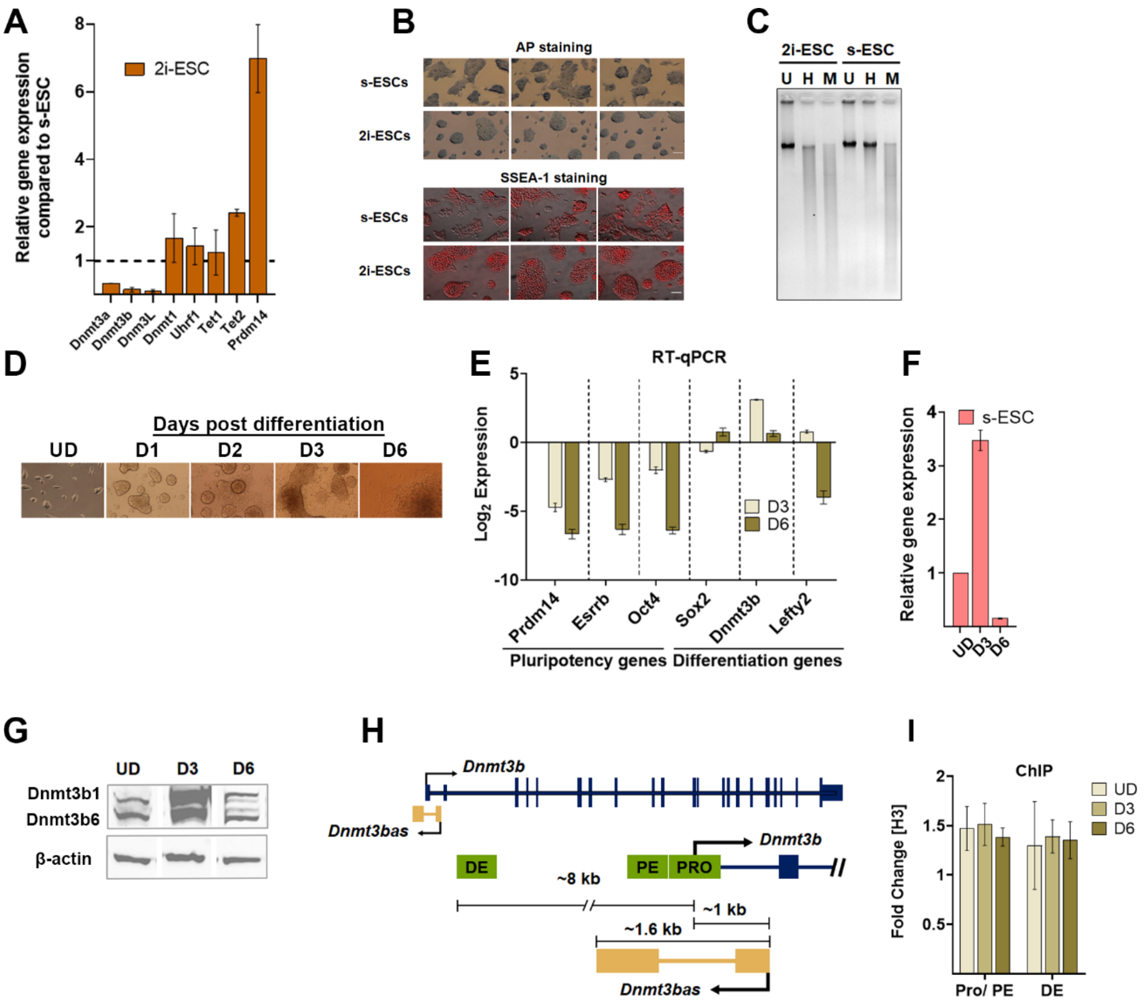
A
Relative gene expression compared to s-ESC
2i-ESC
Dnmt3a
Dnmt3b
Dnm3L
Dnmt1
Uhrf1
Tet1
Tet2
Prdm14
B
AP staining
s-ESCs
2i-ESCs
SSEA-1 staining
s-ESCs
2i-ESCs
C
2i-ESC
s-ESC
U H M U H M
D
Days post differentiation
UD D1 D2 D3 D6
E
RT-qPCR
Log2 Expression
D3
D6
Prdm14
Esrrb
Oct4
Sox2
Dnmt3b
Lefty2
Pluripotency genes
Differentiation genes
F
Relative gene expression
s-ESC
UD
D3
D6
G
UD D3 D6
Dnmt3b1
Dnmt3b6
β-actin
H
Dnmt3b
Dnmt3bas
DE
PE
PRO
Dnmt3b
~8 kb
~1 kb
~1.6 kb
Dnmt3bas
I
ChIP
Fold Change [H3]
UD
D3
D6
Pro/ PE
DE

### Figure S1

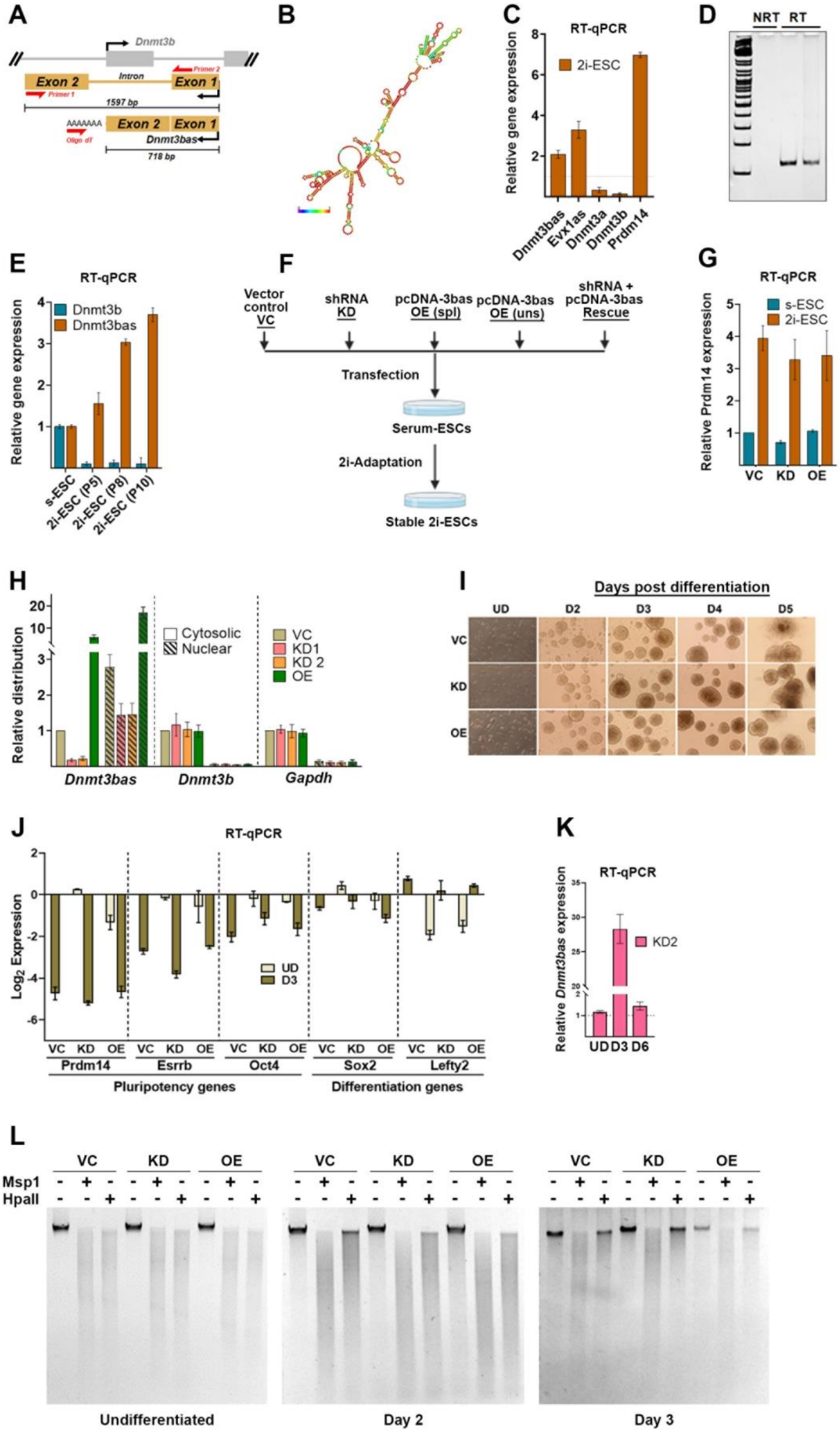

A
Dnmt3b
Exon 2
Intron
Exon 1
Primer 1
Primer 2
1597 bp
AAAAAAA
Oligo dT
Dnmt3bas
718 bp
B
C
RT-qPCR
2i-ESC
Relative gene expression
Dnmt3bas
Evx1as
Dnmt3a
Dnmt3b
Prdm14
D
NRT
RT
E
RT-qPCR
Dnmt3b
Dnmt3bas
Relative gene expression
s-ESC
2i-ESC (P5)
2i-ESC (P8)
2i-ESC (P10)
F
Vector control VC
shRNA KD
pcDNA-3bas OE (spl)
pcDNA-3bas OE (uns)
shRNA + pcDNA-3bas Rescue
Transfection
Serum-ESCs
2i-Adaptation
Stable 2i-ESCs
G
RT-qPCR
s-ESC
2i-ESC
Relative Prdm14 expression
VC
KD
OE
H
Relative distribution
Cytosolic
Nuclear
VC
KD1
KD 2
OE
Dnmt3bas
Dnmt3b
Gapdh
I
Days post differentiation
UD
D2
D3
D4
D5
VC
KD
OE
J
RT-qPCR
Log2 Expression
UD
D3
Prdm14
Esrrb
Oct4
Sox2
Lefty2
Pluripotency genes
Differentiation genes
K
RT-qPCR
Relative Dnmt3bas expression
KD2
UD
D3
D6
L
VC
KD
OE
Msp1
HpaII
Undifferentiated
Day 2
Day 3

**Figure S2**

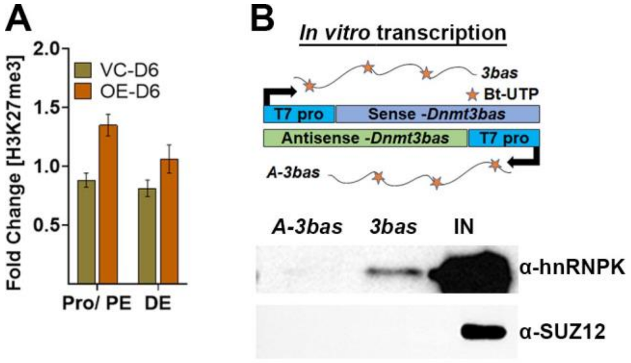
A
Fold Change [H3K27me3]
2.0
1.5
1.0
0.5
VC-D6
OE-D6
Pro/ PE
DE
B
In vitro transcription
3bas
Bt-UTP
T7 pro
Sense -Dnmt3bas
Antisense -Dnmt3bas
T7 pro
A-3bas
A-3bas
3bas
IN
α-hnRNPK
α-SUZ12

**Figure S3**

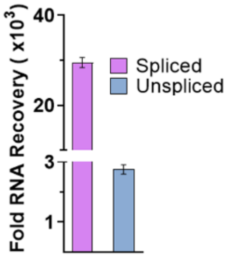

### Figure S4

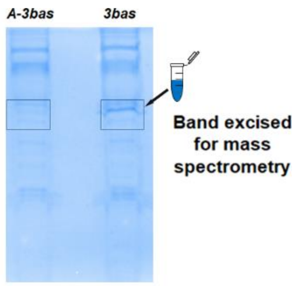

### Figure S5

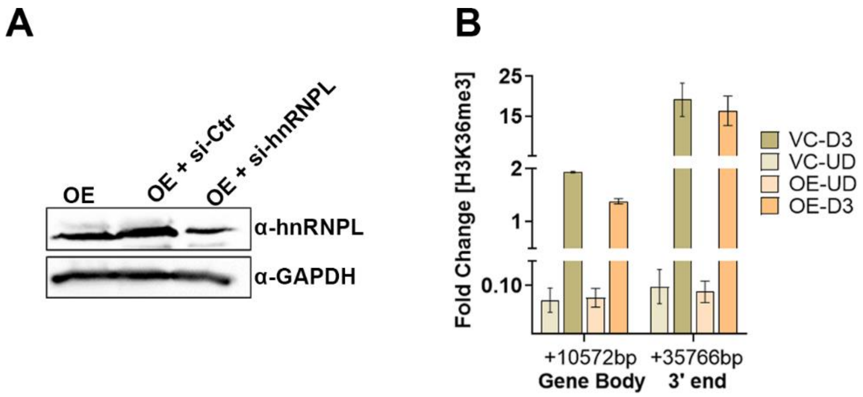
A
OE
OE + si-Ctr
OE + si-hnRNPL
α-hnRNPL
α-GAPDH
B
Fold Change [H3K36me3]
25
15
2
1
0.10
+10572bp
Gene Body
+35766bp
3' end
VC-D3
VC-UD
OE-UD
OE-D3

### Figure S6

#### Supplementary Tables

| For RT-qPCR: | Primer |
|---|---|
| *Dnmt3a RT F* | CCT GCA ATG ACC TCT CCA TT |
| *Dnmt3a RT R* | CAG GAG GCG GTA GAA CTC AA |
| *Dnmt3b RT F* | TGG TGA TTG GTG GAA GCC |
| *Dnmt3b RT R* | AAT GGA CGG TTG TCG CC |
| *Dnmt3L RT F* | ATG GAC AAT CTG CTG CTG ACT G |
| *Dnmt3L RT R* | CGC ATA GCA TTC TGG TAG TCT CTG |
| *Dnmt1 RT F* | GGG TCT CGT TCA GAG CTG |
| *Dnmt1 RT R* | GCA GGA ATT CAT GCA GTA AG |
| *Dnmt3bas RT F* | TCTGGGTAAGAACTGGGCTAGA |
| *Dnmt3bas RT R* | GCAGGGTTGCTGTGTGACT |
| *Dnmt3bas In RT F* | CCTCCCGAGCTTCCTCGCCC |
| *Dnmt3bas E RT R* | GCCGCTCATCCAGACCTGTG |
| *Evx1as-RT F* | CCA GAC ACA GAG GAT GCA AA |
| *Evx1as-RT R* | TGA AGA GCC AAT CCA AAT TCA |
| *Uhrf1 RT F* | GCT CCA GTG CCG TTA AGA CC |
| *Uhrf1 RT R* | CAC GAG CAC GGA CAT TCT TG |
| *Tet1 RT F* | CCA TTC TCA CAA GGA CAT TCA CA |
| *Tet1 RT R* | GCA GGA CGT GGA GTT GTT CA |
| *Tet2 RT F* | GCC ATT CTC AGG AGT CAC TGC |
| *Tet2 RT R* | ACT TCT CGA TTG TCT TCT CTA TTG AGG |
| *Prdm14 RT F* | ACA GCC AAG CAA TTT GCA CTA C |
| *Prdm14 RT R* | TTA CCT GGC ATT TTC ATT GCT C |
| *Sox2 RT F* | ATGCACCGCTACGACGTCAG |
| *Sox2 RT R* | AGGTGAAAGCTTTTATTTTGTTGAGAC |
| *Oct3/4 RT F* | TCT TTC CAC CAG GCC CCC GGC TC |
| *Oct3/4 RT R* | TGC GGG CGG ACA TGG GGA GAT CC |
| *Trim28 RT F* | GAGATGGAGAGCGAACAGTCTAC |
| *Trim28 RT R* | TGTCACAGCTCTCACAGAACAG |
| *Esrrb RT F* | CCTCATCAACTGGGCCAAGC |
| *Esrrb RT R* | TACACGATGCCCAAGATGAGAATCT |
| *Gapdh RT F* | CAAAATGGTGAAGGTCGGTGTGAA |
| *Gapdh RT R* | CAACAATCTCCACTTTGCCACTG |
| *18S RT FP* | AGT CCC TGC CCT TTG TAC ACA |
| *18S RT RP* | GAT CCG AGG GCC TCA CTA AAC |
| *Actb RT F* | TCTTTGCAGCTCCTTCGTTG |
| *Actb RT R* | ACGATGGAGGGGAATACAGC |
| **For ChIP-qPCR:** | |
| *Dnmt3b Pro. F* | TGAAGTCAGGGAGCCAACAA |
| *Dnmt3b Pro. R* | GCAAAGGCTCGGCTCGATA |
| *Dnmt3b DE F* | GCATGAGTGTTCCTGTGGAAA |
| *Dnmt3b DE R* | CTCAGGCTTGATTGTTGTGTCC |
| *Dnmt3b Pro/CGI F* | GCCGCTCATCCAGACCTGTG |
| *Dnmt3b Pro/CGI R* | CCTCCCGAGCTTCCTCGCCC |
| *Gene body Dnmt3b-5F* | GGTGGGAATGCAGGAGCTT |
| *Gene body Dnmt3b-5R* | TGCCATGAAGATTAAAGGCCTAA |
| *3' end Dnmt3b-7F* | TGTGGTGGGATCATGCACAT |
| *3' end Dnmt3b-7F* | AACATCCTGAAACTTTCCTTGTGAA |

| | |
|---|---|
| **For MD-qPCR:** | |
| *Dnmt3b Pro/PE. F* | TGAAGTCAGGGAGCCAACAA |
| *Dnmt3b Pro/PE. R* | GCAAAGGCTCGGCTCGATA |
| *Dnmt3b DE F* | GCATGAGTGTTCCTGTGGAAA |
| *Dnmt3b DE R* | CTCAGGCTTGATTGTTGTGTCC |
| *-329 Dnmt3b Pro. F* | GCCGCTCATCCAGACCTGTG |
| *-329 Dnmt3b Pro. R* | CCTCCCGAGCTTCCTCGCCC |
| *-373 Dnmt3b Pro. F* | GCAGGGTTGCTGTGTGACT |
| *-373 Dnmt3b Pro. R* | GCACACACGCACATACAAGC |
| *-434 Dnmt3b Pro. F* | CCATAGGGCCACTACAGCGC |
| *-434 Dnmt3b Pro. R* | GCACACACGCACATACAAGC |
| *-861 Dnmt3b Pro. F* | TGAAGTCAGGGAGCCAACAA |
| *-861 Dnmt3b Pro. R* | GCAAAGGCTCGGCTCGATA |
| *Dnmt3b Pro/CGI F* | GCCGCTCATCCAGACCTGTG |
| *Dnmt3b Pro/CGI R* | CCTCCCGAGCTTCCTCGCCC |
| *Ctrl F* | ACCTAAACCTCATAAAGACACAACA |
| *Ctrl R* | TGACGTGTTCTTGGATTCAGT |
| *H19 F* | CCG TTT TAG GAC TGC GAT GT |
| *H19 R* | GGG TCA CAA ATG CCA CTA GG |
| **For 3C:** | |
| *Dnmt3b 3C F1 (anchor)* | ACCAAATTCCAGGTCAGTCTGG |
| *Dnmt3b 3C R3* | GGATGTACGGGGGTCTGTCA |
| *Dnmt3b 3C R4* | TTCTCTTACCTCGGCTGGGA |
| *Dnmt3b 3C R8* | GCATCCTCTTTTGCACCAACT |
| *Dnmt3b 3C R9* | GTGAAGTCAGGGAGCCAACA |
| *Dnmt3b 3C R10* | GCACGAGGAACCCAGGTAG |
| *Dnmt3b 3C R13* | ACCTCTAGTTCTGGATACCATTTCC |
| **For Cloning:** | |
| *Dnmt3bas-Exon1-FP* | GTGTGGTGGAATTCTGCAGATAGCCCCTCCCAGC |
| *Dnmt3bas-Exon1-RP* | GTGTGCGCGGCCGCTAGCCCAGTTCTTACCCAGAGCCTCCTCGGGGAGG |
| *Dnmt3bas-Exon2-FP* | GAACTGGAATTCATCAGATGCACACACGCACATACAAGCACATTCAC |
| *Dnmt3bas-Exon2-RP* | GACTCGAGCGGCCGCCACTGTGCTGGATCCCTTTAAC |
| *pLKO.1 sequencing primer* | CAA GGC TGT TAG AGA GAT AAT TGG A |
| **For Bisulfite Seq.** | |
| *Cxxc5 F* | AGTAGTAGTAATATTAATAGTAGTAGTG |
| *Cxxc5 R* | ACTCCTTATTAAAAAATTCACTAATAATAC |
| *Rbm12b2 F* | AAAAGGGAATAAAGAAGAAAAAGAATTA |
| *Rbm12b2 R* | ATAACTTACCACTAAATAAACAAACAAA |

**Table S1. Primers used in this study.**

A list of all PCR primers used in this study (5’ to 3’), separated by technique.

| *Sequence Name* | *Sequence* |
|---|---|
| *Dnmt3bas-Chrip_1* | TTGTGCCAGACCTTGGAA |
| *Dnmt3bas-Chrip_2* | AGAGTGCTTCCGGACTTG |
| *Dnmt3bas-Chrip_3* | AGTCCTGTGATCTCCATG |
| *Dnmt3bas-Chrip_4* | ACCTACTTACTGCGTTCC |
| *Dnmt3bas-Chrip_5* | GGTTAATCGTCCGTGCTT |
| *Dnmt3bas-Chrip_6* | ATTCTTAGCCTTCTTCCG |
| *Dnmt3bas-Chrip_7* | CTGGCTGAGCACACCTGG |
| *Dnmt3bas-Chrip_8* | ATGCTAGATGGGCGTTGT |
| *Dnmt3bas-Chrip_9* | GTAGTGGTTGTGGATCCG |
| *Dnmt3bas-Chrip_10* | ACACTTGGGAGCGGAGTG |
| *Dnmt3bas-Chrip_11* | CAGAGGAAGCGAAGTCCC |
| *Dnmt3bas-Chrip_12* | GTGGCGCCTGGATGTTTG |
| *Dnmt3bas-Chrip_13* | AGCCTCACGACAGGTGAG |
| *Dnmt3bas-Chrip_14* | GCGGCCCAAGTAAACGTA |
| *Dnmt3bas-Chrip_15* | TGGACGCTCTCGCCTGAG |
| *Dnmt3bas-Chrip_16* | CGGGCTACAAGGGCAGCG |
| *Dnmt3bas-Chrip_17* | GCTCGGGAGGGATTTCAA |
| *Dnmt3bas-Chrip_18* | TGGGCTCTGGTCATCTAG |
| *Dnmt3bas-Chrip_19* | CCAGACCTGTGAATGTGC |
| *Dnmt3bas-Chrip_20* | AGCGGAGGGATGGTCGAA |
| *Dnmt3bas-Chrip_21* | ACAAAGGCCAAGCTAGGT |
| *Dnmt3bas-Chrip_22* | GCACGAGGAACCCAGGTA |
| *Dnmt3bas-Chrip_23* | TGTGTTTCTCCAGTGGTT |
| *Dnmt3bas-Chrip_24* | GCTAGGCCTTGTATCGAG |
| *Dnmt3bas-Chrip_25* | GCTTGGTGTGTAGGAGGA |
| *Dnmt3bas-Chrip_26* | GACTGTTCTGTTTGGTGA |
| *Dnmt3bas-Chrip_27* | ACCTGTGCTCCTAGAGAG |
| *Dnmt3bas-Chrip_28* | GAACCCCAGAGGCAAGAG |
| *Dnmt3bas-Chrip_29* | CCCAAATAGTCGCCATTT |

**Table S2. ChIRP probes used in this study.**

A list of all ChIRP probes used in this study (5’ to 3’) with 3' biotin modification

### Supplementary Materials and Methods

#### In-Gel Trypsin digestion and Mass Spectrometry

The RNA-pull down protein eluates were resolved with SDS-PAGE, followed by Coomassie staining. The observed unique 60 kDa band in the *Dnmt3bas* pulldown lane, which was absent in anti-sense *Dnmt3bas* lane was excised together with the corresponding region in the control lane. Proteins in the excised gel were destained and in-gel trypsin digested to obtain peptides as previously described (Shevchenko et al., 2006). The peptides were reconstituted in 50% acetonitrile/0.1 % TFA and used for MALDI-TOF mass spectrometry to identify potential *Dnmt3bas* binding partners.

#### *Dnmt3bas* secondary structure determination

The nucleotide sequence of spliced *Dnmt3bas* was used to predict secondary structures using the default fold algorithm and basic options on the Vienna RNA fold webserver (http://rna.tbi.univie.ac.at/#databases).

#### Microscopy

Bright field images of s-ESCs and 2i-ESCs were obtained with Zeiss microscope using a 10X 591 objective. We performed Alkaline phosphatase staining and SSEA-1 staining in these cells as previously reported (AlAbdi *et al.*, 2020). Briefly, the cells were stained with alkaline phosphatase (Sigma, AB0300) and SSEA-1 staining using (Millipore, 597 MAB430) and AlexaFluor 555 nm (Life Technologies, A21422) antibodies. The SSEA-1 and Alkaline 598 phosphatase stained cells were imaged using 20X objectives under Nikon Ts and Zeiss 599 microscopes, respectively.

#### Genome-wide DNA methylation Profile

Genomic DNA was isolated using the standard phenol-chloroform method. Following RNase digestion and re-purification by phenol-chloroform, 1 ug of DNA was digested with either HpaII or MspI overnight at 37 $^{o}$C. As a control, 1 ug of DNA was incubated with digestion buffer under same conditions. Undigested and digested DNA samples were run on a 1% agarose Tris-acetic acid EDTA gel for 90 mins at a constant 100 V. Following ethidium bromide staining, the gel was imaged with Axygen® gel documentation system (Corning, GD-1000).

#### Supplementary References

AlAbdi, L., Saha, D., He, M., Dar, M.S., Utturkar, S.M., Sudyanti, P.A., McCune, S., Spears, B.H., Breedlove, J.A., Lanman, N.A., and Gowher, H. (2020). Oct4-Mediated Inhibition of Lsd1 Activity Promotes the Active and Primed State of Pluripotency Enhancers. Cell Reports *30*, 1478-1490.e1476. https://doi.org/10.1016/j.celrep.2019.11.040.

Shevchenko, A., Tomas, H., Havlis, J., Olsen, J.V., and Mann, M. (2006). In-gel digestion for mass spectrometric characterization of proteins and proteomes. Nat Protoc *1*, 2856-2860. 10.1038/nprot.2006.468.
